# Intestinal helminth skews DC2 development towards regulatory phenotype to counter the anti-helminth immune response

**DOI:** 10.1101/2024.09.11.612410

**Authors:** Anna Andrusaite, Olivia Ridgewell, Anna Ahlback, Holly Webster, Hiroki Yamaguchi, Molly Peel, Annika Frede, Sarwah Al-Khalidi, Andrew Farthing, Anna Heawood, Annabelle Smith, Edward Roberts, Allan Mowat, Richard Maizels, Georgia Perona-Wright, Simon Milling

## Abstract

The intestinal immune system maintains a balance between active immunity needed for protection and tolerance towards harmless antigens. Dendritic cells (DCs) found in the intestinal mucosa are key to the adaptive arm of these immunoregulatory events. DCs sample antigens in the tissue and then migrate to the draining lymph nodes, where they prime the T cells that then migrate back to the tissue as effector or regulatory cells. Intestinal DC are highly heterogeneous, and it remains unclear exactly which subsets induces the different kinds of immune response, or what signalling molecules and cellular mechanisms are involved. Here, we have studied these issues using *Heligmosomoides polygyrus bakeri (Hpb)* infection in mice, a model which is uniquely suited to dissecting this regulatory circuit in the gut, where it drives type 2 protective immunity at the same time as inhibiting other aspects of the immune response. Here, we characterise intestinal DC during *Hpb* infection for the first time. We observed a dynamical change of intestinal DC populations throughout the course of infection that correlated with altered phenotype and function. In particular, *Hpb* infection saw a rise in a population of CD103^+^ DC2 that retained a potent ability to drive Tregs during the infection and unlike CD103-DC2, had a reduced ability to induce pro-inflammatory immune response. Furthermore, transcriptional analysis revealed that TGFβ signalling may be responsible for some of the changes observed. This was confirmed *in vitro*, where supplementation TGFβ or *Hpb*-produced TGFβ mimic (TGM) replicated the immunomodulatory effects seen in DCs *in vivo*. Together, these results present a mechanistic explanation of how helminths such as *Hpb* may modulate host immune responses by altering the differentiation and function of local DCs. Furthermore, our work provides the basis for understanding immune homeostasis in the intestine at the molecular and cellular levels. Thus, this work fills out a crucial gap in our knowledge of basic biology underlining the DC decision between pro- and anti-inflammatory immune response in the central circuit of adaptive immune response.

## Introduction

While *Heligmosomoides polygyrus bakeri* (*Hpb*) has been extensively used as a helminth and Type 2 immunity model for the past couple of decades ^1–5^, surprisingly there has been minimal work done exploring immune cells in the primary site of infection – the small intestine (SI) lamina propria (LP). Physiological changes of the epithelium and LP as well as increased mucus production during infection contribute to the difficulty in isolating viable single cells from these tissues. Previously, we have solved this long-standing issue by development of novel protocol for cell isolation during *Hpb* infection^6^. Furthermore, we used this protocol and identified changes in T cell populations that participate in the mounting of the appropriate immune response^7^. Now, we seek to use this protocol in order to characterise intestinal dendritic cells (DC), which are the key cell driving the adaptive immune response and instruct and control the T cell response. The current understanding of which DC initiate and regulate Th2 immune responses in the gut in response to *Hpb* is very limited and mostly associated with cells isolated from the draining lymph nodes^8^. In other models of Type 2 immune repones, DC2 have been implicated in the driving of Type 2 T helper cells (Th2)^9–11^, however there remains to be population-specific ambiguity and differences between SI and colon^12,13^. Furthermore, same subsets have been shown to also drive other types of immune response as well as induction of T regulatory (Treg) cells in a context dependant manner^14^. Thus, there is an overall lack of clarity around what molecular pathways determine DC ability to drive Th2 cells vs. Treg and the underlying mechanisms remain to be determined.

Given the growing evidence of *Hpb* immunomodulatory abilities^15^, it also remains to be shown if and how intestinal DC may be affected and what it means for the overall adaptive immune response.

### Results

### Effects of *Hp* infection on the intestinal DC populations

To explore how DC populations behaved during *Hpb* infection, we used our recently published method^6^ to isolate these cells from small intestine lamina propria (SI LP) over the first 14 days of infection, during which time *Hpb* breeches the intestinal barrier twice and in-between establishes itself in the gut wall for a mandatory maturation stage (as summarised in Figure S1a). Isolated DC were subdivided using CD103 and CD11b as previously described^14^ (Figure S2b). Use of these two markers allows identification of DC1 (CD103^+^ CD11b^-^), two distinct DC2 populations defined by their CD103 expression (CD11b^+^CD103^-^ and CD11b^+^CD103^+^ DC2) and double negative (DN) cells, that are believed to be a mixed pool of immature DC with minimal immune potential and poorly characterised mature DC. We quantified SI LP DC population proportional distribution throughout the infection and identified an increase in CD103^+^ DC2 around 5 days post infection (5 DPI), that peaks at 9 DPI and returns to naïve levels after that (Figure S1c). This increase led a proportional decrease of CD103^-^ DC2 and DC1, however, when absolute numbers of cells were quantified, along with the increase in CD103^+^ DC2 abundance, other cells also increase between 5 DPI-9 DPI, and then subsequently decreased (Figure S1d). As the peak of changes in SI LP DC populations occurred around 7DPI and this timepoint is widely considered to be the peak of the inflammatory response in this infection model, we decided to further explore SI LP DCs behaviour in more detail. First, we analysed SI LP DC populations at 7DPI (Figure 1a). This confirmed a significant increase in the proportion of CD103^+^ DC2 in *Hpb* infected mice compared with uninfected naïve mice (Figure 1b). Although the proportion of CD103-DC2 decreased in parallel, the absolute number of both DC2 populations was significantly increased in infected mice (Figure 1c). The numbers of DC1 also increased significantly during *Hpb* infection, but their proportions were unchanged compared with controls. Given that DC2 are largely responsible for the protective Type 2 immunity during helminth infection, we focused the rest of the study on populations within the DC2 lineage.

**Figure 1.**
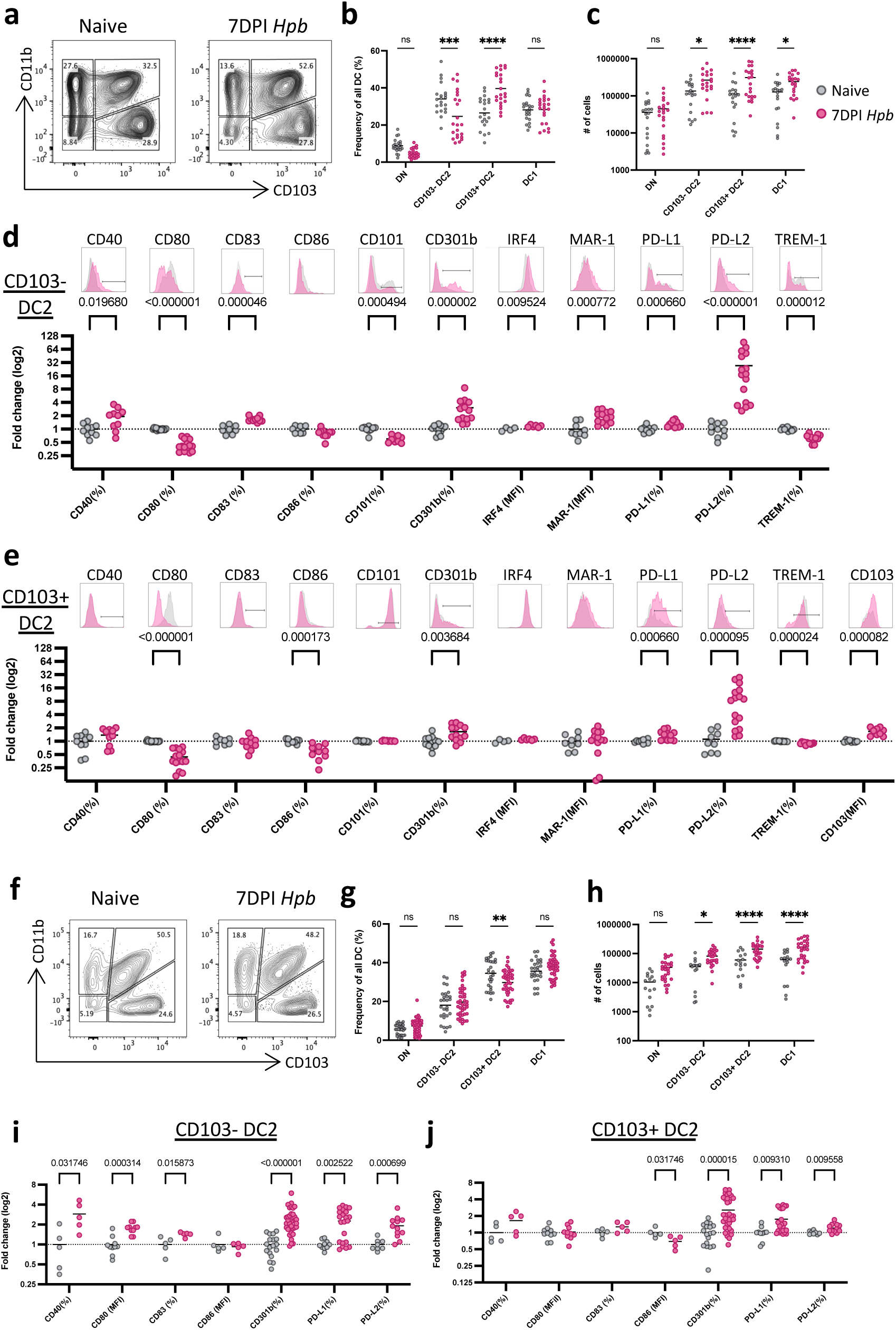
Changes in phenotypically distinct DC populations at 7DPI of *Hpb* infection in the SI LP and sMLN. A) Representative contour plots of DC subsets based on their CD103 and CD11b expression. Isolated from SI LP at 7DPI with *Hpb* or naïve controls. DCs were identified as live, single, CD45^+^, CD3^-^, B220^-^, CD64^-^, MHCII^+^ and CD11c^+^. B) Quantified frequency of DC subsets isolated from SI LP at 7DPI with *Hpb* or naïve controls. Data shown are pooled from 6 independent experiments (n=3-5 each) C) Quantified absolute cell number of DC subsets isolated from SI LP at 7DPI with *Hpb* or naïve controls. Data shown are pooled from 6 independent experiments (n=3-5 each). D) Fold change of phenotypical markers in CD103^-^ DC2 isolated from SI LP at 7DPI with *Hpb* or naïve controls. Data shown are pooled from 2-3 independent experiments for each marker (n=3-5 each). E) Fold change of phenotypical markers in CD103^+^ DC2 isolated from SI LP at 7DPI with *Hpb* or naïve controls. Data shown are pooled from 2-3 independent experiments for each marker (n=3-5 each). F) Representative contour plots of migratory DC subsets based on their CD103 and CD11b expression. Isolated from sMLN at 7DPI with *Hpb* or naïve controls. DCs were identified as live, single, CD45^+^, CD3^-^, B220^-^, CD64^-^ , MHCII^hi^ and CD11c^+^. G) Quantified frequency of migratory DC subsets isolated from sMLN at 7DPI with *Hpb* or naïve controls. Data shown are pooled from 6 independent experiments (n=3-5 each) H) Quantified absolute cell number of migratory DC subsets isolated from sMLN at 7DPI with *Hpb* or naïve controls. Data shown are pooled from 6 independent experiments (n=3-5 each). I) Fold change of phenotypical markers in migratory CD103^-^ DC2 isolated from sMLN at 7DPI with *Hpb* or naïve controls. Data shown are pooled from 1-5 independent experiments for each marker (n=5-7 each). J) Fold change of phenotypical markers in CD103^+^ DC2 isolated from SI LP at 7DPI with *Hpb* or naïve controls. Data shown are pooled from 1-5 independent experiments for each marker (n=5-7 each). p-values shown (D, E, I, J) or represented as *p<0.05, **p<0.01, ***p<0.001,****p<0.0001 as assessed by two-way ANOVA with Šídák’s post-test correction for multiple comparisons. ns=not significant.

Next, we sought to determine possible functional implications in the changes observed in the DC2 compartment during *Hpb* infection. First, we characterised the DC phenotype based on markers that can associated with priming and shaping of adaptive immune responses (CD80, CD83, CD86, MAR-1)^16–18^, molecules previously associated with DC2 ability to induce Th2 response (CD40, CD301b, PD-L2, IRF4)^13,19–22^ and molecules associated with DC regulatory function (CD101, PD-L1, TREM-1)^23–26^ (Figure 1d,e). We identified that CD103^-^ DC2 isolated from *Hpb* infected SI LP 7DPI have an increased proportion of CD40^+^, CD83^+^, CD301b^+^, PD-L1^+^ and PD-L2^+^ cells and these cells also had an overall increased levels of IRF4 and MAR-1 measured as an MFI (mean fluorescence intensity). However, it was also observed that there was a decreased proportion of CD80^+^, CD101^+^ and TREM-1^+^ cells while CD86^+^ cells proportion did not change (Figure 1d). Together, this phenotypical landscape suggests a pro-inflammatory and Th2-skewing signature. On contrary, while CD103^+^ DC2 had a similarly increased fold change in proportion of CD301b^+^, PD-L1^+^, PD-L2^+^ cells, there was no increased observed in context of CD40, CD83, IRF4 and MAR-1 (Figure 1e). Similar to CD103^-^ DC2, within the CD103^+^ DC2 populations there was a decreased proportion of CD80^+^ cells, a small decrease in TREM-1^+^ cells, as well as additionally decreased proportion of CD86^+^ cells. IRF4 and MAR-1 expression pattern did not change. Furthermore, CD103^+^ DC2 maintained a high level of CD101 and increased overall expression levels of CD103, as measured by MFI. Together, CD103^+^ DC2 phenotypical landscape was different from the CD103^-^ DC2. Furthermore, the distinct phenotype of CD103^+^ DC2 indicate a reduced ability for driving the pro-inflammatory and Th2 immune response, and instead suggest a possible regulatory function.

Given the previously observed changes of DC abundance throughout the *Hpb* infection (Figure S1b-d) and the known biology of *Hpb* lifecycle that involves multiple barrier breeches as well as establishment in the SI gut wall^27^ (Figure S1a), we sought to establish if there is a correlation between the phenotypical changes of DC2 and timepoint of the infection. We selected to assess CD40, CD80, CD83, CD301b and PDL2 (Figure S1e-i) as these markers previously showed the most significant change at 7DPI in DC2 populations (Figure 1d, e). The proportion of CD40^+^ cells increased within the CD103^-^ DC2 population around 5 DPI and was sustained until 12 DPI, when it returned to naïve levels. This increase was significantly higher than the almost unchanged proportion of CD40^+^ cells within CD103^+^ DC2 (Figure S1e). The proportion of CD80^+^ cells steadily decreased in CD103^+^ DC2 population, while it was somewhat varied within the CD103^-^ DC2 population (Figure S1f). The proportion of CD83^+^ cell increased within both DC2 populations early in the infection, after the first barrier breech around 2PDI. This increase was sustained in the CD103^-^ DC2 population throughout the course of the infection, while stunted within the CD103^+^ DC2 population (Figure 1Sg). The proportion of CD301b^+^ cells increased steadily in both DC2 populations and interestingly, was higher within the CD103^+^ DC2 around 12-14DPI in comparison to the CD103-DC2 counterparts (Figure S1h). Finally, while both DC2 subsets had an increased proportion of PD-L2+ cells after 2DPI and then again around 9DPI, this proportion was significantly higher within the CD103^-^ DC2 population between 9-12 DPI (Figure S1i). Taken together, this data illustrates that both, the SI LP DC2 abundance and phenotype dynamically change through-out the *Hpb* infection and often correlate with known timepoints in *Hpb* life cycle, such as barrier breeches at 2DPI and 9DPI or the helminth establishment in the gut wall between these two time points. Additionally, the 7DPI timepoint was identified to be peak of these changes, thus we used it in our further analysis. Finally, our observations imply that *Hpb* presence during the infection may shape the intestinal environment and affect intestinal DC abundance and phenotype.

Upon antigen uptake DC leave the tissue and migrate to the intestinal draining lymph nodes where they interact with naïve T cells to drive adaptive immune response. Therefore, to explore whether the changes in DC subsets and phenotypes we had seen in SI LP might be relevant to the priming environment, we analysed migratory DC subsets in the small intestine draining mesenteric lymph nodes (sMLN) on 7DPI of *Hpb* infection (Figure1f-j). Here, we identified that contrary to the SI LP, there is a small decrease in CD103^+^ DC2 proportion (Figure 1g). However, the total numbers of all three subsets increased significantly in sMLN isolated during *Hpb* infection (Figure 1h). Thus, the proportional decrease of CD103^+^ DC2 may represent an artefact due to the large cell number increases in CD103^-^ DC2 and DC1. Next, we characterised the phenotypical landscape of DC2 and identified that similarly to the SI LP, migratory CD103^-^ DC2 isolated from the infected sMLN had an increase proportion of CD40^+^, CD80^+^, CD83^+^, CD301b^+^, PD-L1^+^ and PD-L2^+^ cells (Figure 1i), while within the migratory CD103^+^ DC2 population, the proportion of CD40^+^, CD80^+^ and CD83^+^ cells did not change. Yet, a decrease in the CD86^+^ cell proportion was observed as well as an increase in CD301b^+^, PD-L1^+^ and PD-L2^+^. Taken together, the phenotypical landscapes of migratory DC2 appear to partially mirror the phenotype of their counterparts in the SI LP during *Hpb* infection. Additionally, the phenotypical difference between the two DC2 populations found in SI LP is also partly conserved in the migratory populations and could therefore suggest a functional divergence of the two DC2 populations during *Hpb* infection.

Together, these findings indicate that Hp infection drives phenotypic changes within mucosal DC2 populations, but the pattern differs between the subsets defined by expression of CD103. In particular, CD103^-^ DC2 appear to acquire more of a pro-inflammatory phenotypic landscape than their CD103^+^ counterparts and this may show somewhat of bias toward a Th2-skewing signature. In contrast, the changes in the CD103^+^ DC2 population seem to favour more of an anti-inflammatory phenotype. Having established distinct phenotypical landscapes within the two DC populations in the SI LP, we next sought to understand if these changes also translate into altered function of the intestinal DC2.

### SI LP DC function is altered during *Hp* in a subset specific manner

Here we examined the possible functional consequences of the phenotypic changes we had observed in DC2 in response to *Hpb* infection. To test this, we FACS sorted SI LP DC populations from naïve and *Hpb* infected SI LP at 7DPI, pulsed them with ovalbumin protein (OVA) and co-cultured them with CTV-labelled, naïve, OVA-specific TcR transgenic CD4^+^ T cells. Under these conditions, all subsets SI LP DCs were able to drive T cell proliferation, as measured by CTV dilution (Figure 2a). This ability was increased in CD103^+^ DC2 and DC1 isolated from the infected intestine (Figure 2b). Similarly, the same change was observed when cultured T cells were assessed based on their CD44 and CD69 expression, that represents T cell activation (Figure 2c-f) with CD103^+^ DC2 and DC1 isolated from the infected intestine increased in their ability to drive T cells expression of CD44 (Figure 2d), while all subsets had an increased ability to stimulate CD69 expression (Figure 2f). Next, as the DC phenotypical changes during *Hpb* infection suggested a possible regulatory function, we analysed T cells for Foxp3 expression which is used to identify regulatory T cells (Tregs) (Figure 2g). Here we identified that while within the DC isolated from naïve SI LP DC1 are the most prominent drivers of Tregs, during *Hpb* infection this ability is lost; and yet, CD103^+^ DC2 isolated from Hp SI LP retained their ability to induce Tregs (Figure 2h). Additionally, we also analysed cytokines present in these co-cultures (Figure 2i). Here, we identified that CD103^-^ DC2 isolated from *Hp* SI LP had an increased ability to stimulate T cells to produce IL-2, IL-6, IL-10, IL-13, IL-17A, INFγ and TNFα. However, CD103^+^ DC2 isolated from the infected tissue had an increased ability to stimulate T cell production of IL-2 and TNFα, but IL-6, IL-10, IL-13, IL-17A and INFγ levels were unchanged and IL-4 levels were decreased.

**Figure 2.**
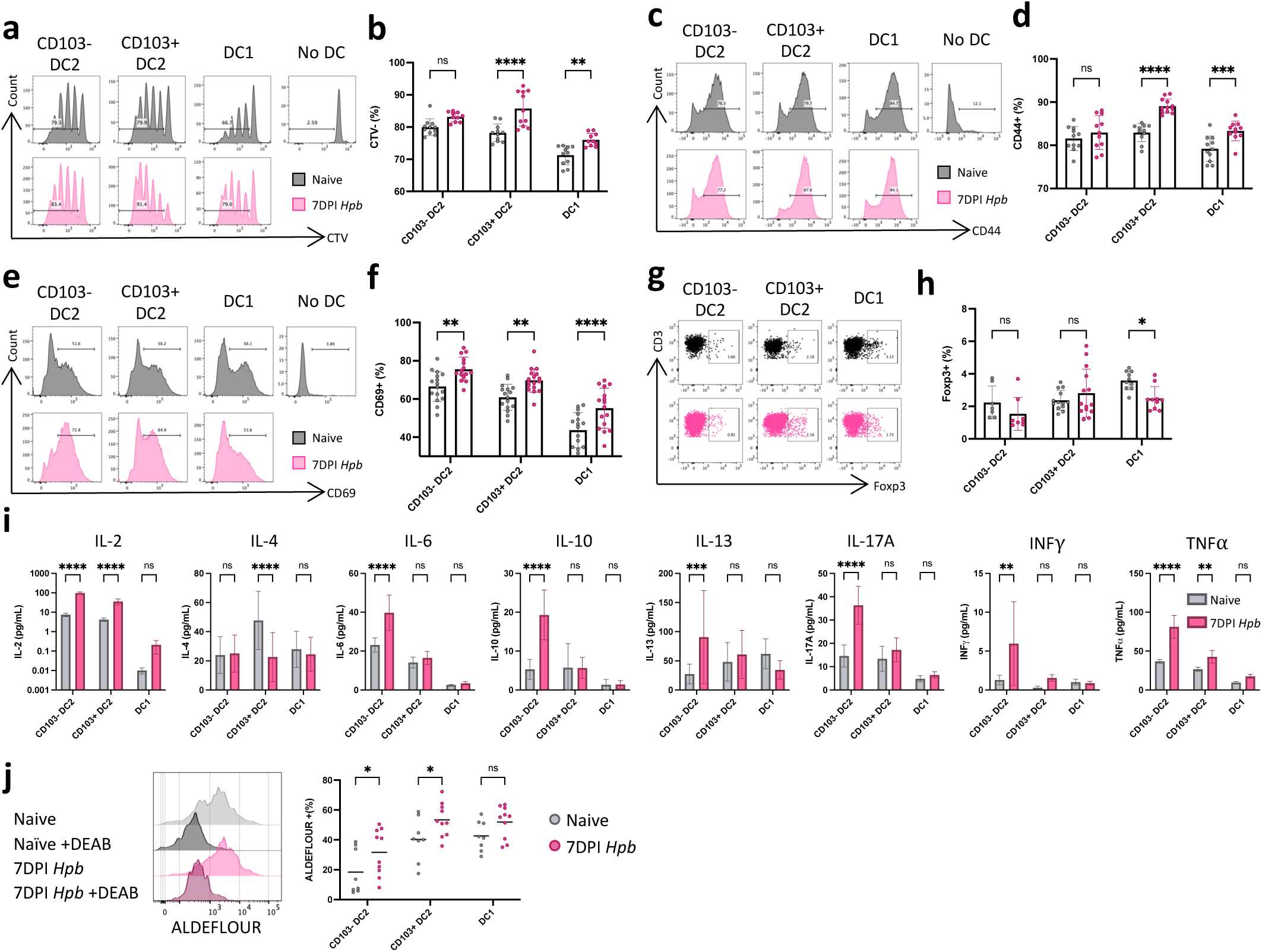
SI LP DC2 function is altered in subset-specific manner at 7DPI with *Hp*. A) Histograms showing levels of cell trace violet (CTV) expression by naïve OTII CD4^+^ T cells cultured at a 10:1 ratio for 3 days with OVA-pulsed DC subsets isolated SI LP at 7DPI with *Hpb* or naïve controls. B) Quantified proportions of CTV^-^ within OT-II CD4^+^ T cells after 3 days of co-culture with OVA-pulsed DC subsets isolated SI LP at 7DPI with *Hpb* or naïve controls. Data shown are pooled from 2 independent experiments (n=5-6 each). C) Histograms showing levels of CD44 expression by naïve OTII CD4^+^ T cells cultured at a 10:1 ratio for 3 days with OVA-pulsed DC subsets isolated SI LP at 7DPI with *Hpb* or naïve controls. D) Quantified proportions of CD44^+^ within OT-II CD4^+^ T cells after 3 days of co-culture with OVA-pulsed DC subsets isolated SI LP at 7DPI with *Hpb* or naïve controls. Data shown are pooled from 2 independent experiments (n=5-6 each). E) Histograms showing levels of CD69 expression by naïve OTII CD4^+^ T cells cultured at a 10:1 ratio for 3 days with OVA-pulsed DC subsets isolated SI LP at 7DPI with *Hpb* or naïve controls. F) Quantified proportions of CD69^+^ within OT-II CD4^+^ T cells after 3 days of co-culture with OVA-pulsed DC subsets isolated SI LP at 7DPI with *Hpb* or naïve controls. Data shown are pooled from 2 independent experiments (n=5-6 each). G) Dotplot showing proportion of Foxp3 expression by naïve OTII CD4^+^ T cells cultured at a 10:1 ratio for 3 days with OVA-pulsed DC subsets isolated SI LP at 7DPI with *Hpb* or naïve controls. F) Quantified proportions of Foxp3^+^ within OT-II CD4^+^ T cells after 3 days of co-culture with OVA-pulsed DC subsets isolated SI LP at 7DPI with *Hpb* or naïve controls. Data shown are pooled from 2-3 independent experiments (n=3-5 each). I) Quantification of cytokines in the supernatant of naïve OTII CD4^+^ T cells cultured at a 10:1 ratio for 3 days with OVA-pulsed DC subsets isolated SI LP at 7DPI with *Hpb* or naïve controls. Data shown are pooled from 1-3 independent experiments (n=5-6 each). J) Representative histograms (left) and quantification (right) of ALDEFLOUR^+^ cells within DC subsets isolated from SI LP at 7DPI with *Hpb* or naïve controls. Data shown are pooled from 2 independent experiments (n=5 each). *p<0.05, **p<0.01, ***p<0.001,****p<0.0001 as assessed by two-way ANOVA with Šídák’s post-test correction for multiple comparisons. ns=not significant.

Production of retinoic acid (RA) is a characteristic property of intestinal DC and is important for their ability to induce Treg cell differentiation^28,29^. Therefore, we next tested the ability of DC subsets for activity of the aldehyde dehydrogenase (ALDH) enzyme needed to produce RA, using the ALDEFLUOR™ assay. All DC subsets from control SI LP could induce some production of RA, with CD103-DC2 and DC1 having significantly more activity than the CD103^-^ DC2 subset (Figure 2j). RA production by both subsets of DC2 was significantly increased during *Hpb* infection, with this effect being most marked for the CD103^-^ subset. In contrast DC1 ALDH activity was not significantly altered.

Together these results indicate that *Hpb* infection alters the functional profile of SI LP DC2. In particular and consistent with the phenotypic changes we observed, CD103^-^ DC2 from infected mice appear to be more activated and to have an overall pro-inflammatory profile. In contrast, although CD103^+^ DC2 induced more T cell activation and RA production, this subset retained a generally neutral profile and Treg driving ability, which is consistent with their phenotypic changes. Thus, infection with *Hpb* leads to dramatically distinct effects on the two DC2 populations in SI LP, suggesting they may contribute to opposite arms of the immune response against the helminth. As in other models both DC2 populations have been reported to be able to drive anti-helminth Th2 immune response^12^, it remains unclear if in the case of *Hpb*, CD103^-^ and CD103^+^ DC2 have a differently ability to response to the worm antigens or if the inflammation environment is skewing the DC development and activity.

### *Hp* derived antigens are trafficked and processed differently between the two DC2 populations

First, we tested the DC ability to process and present the helminth antigens using excretory/secretory material isolated from *Hpb* (HES)^30^. First, we set out to assess antigen trafficking to the lymph nodes. As analysis of sMLN DC is often flawed due to the short DC lifespan upon arrival in the node, we used a method of surgically acquiring afferent lymph as previously described^31,32^ and illustrated in Figure S2a. This method allowed us to identify migratory DC before their entrance in the nodes, thus avoiding the subsequent changes they might go through in the node, such as induction of apoptosis.

The composition of the DC populations in steady state draining lymph mirrored that in the SI LP, with CD103^+^ DC2 and DC1 being the most abundant populations (30-40%), while CD103^-^ DC2 and DN DC represented smaller proportions (10-20%) (Figure 3a). Next, to test antigen carrying ability under steady state conditions we used a fluorescently conjugated OVA that was injected in the SI LP 16hrs prior to the lymph collection (Figure S2a). We then isolated DC from the afferent lymph and quantified OVA^+^ antigen carrying DCs (Figure S2b). We identified that OVA^+^ DC distribution did not mirror lymph DC subset frequencies identified in Figure 3a. Instead, the majority (40%) of OVA trafficking DC were CD103^-^ DC2, then followed by DC1 (30%), while the proportion CD103+ DC2 carrying OVA was much smaller (20%) (Figure 3b). Using the same approach we assessed SI LP DC ability to carry *Hpb* antigens using fluorescently conjugated HES (Figure S2c). In contrast to the DC distribution frequencies of the OVA-trafficking DCs, CD103^-^ DC2 appeared to be predominantly carrying HES in the afferent lymph (50%), followed by CD103+ DC2 (20-30%) and much smaller population of DC1 (10-20%) (Figure 3c). Thus, unlike OVA, HES antigens are preferentially carried in lymph by DC2, predominantly by CD103^-^ DC2 subset. Thus, HES antigens are preferentially carried in lymph by DC2, predominantly by the CD103^-^ DC2 population.

**Figure 3.**
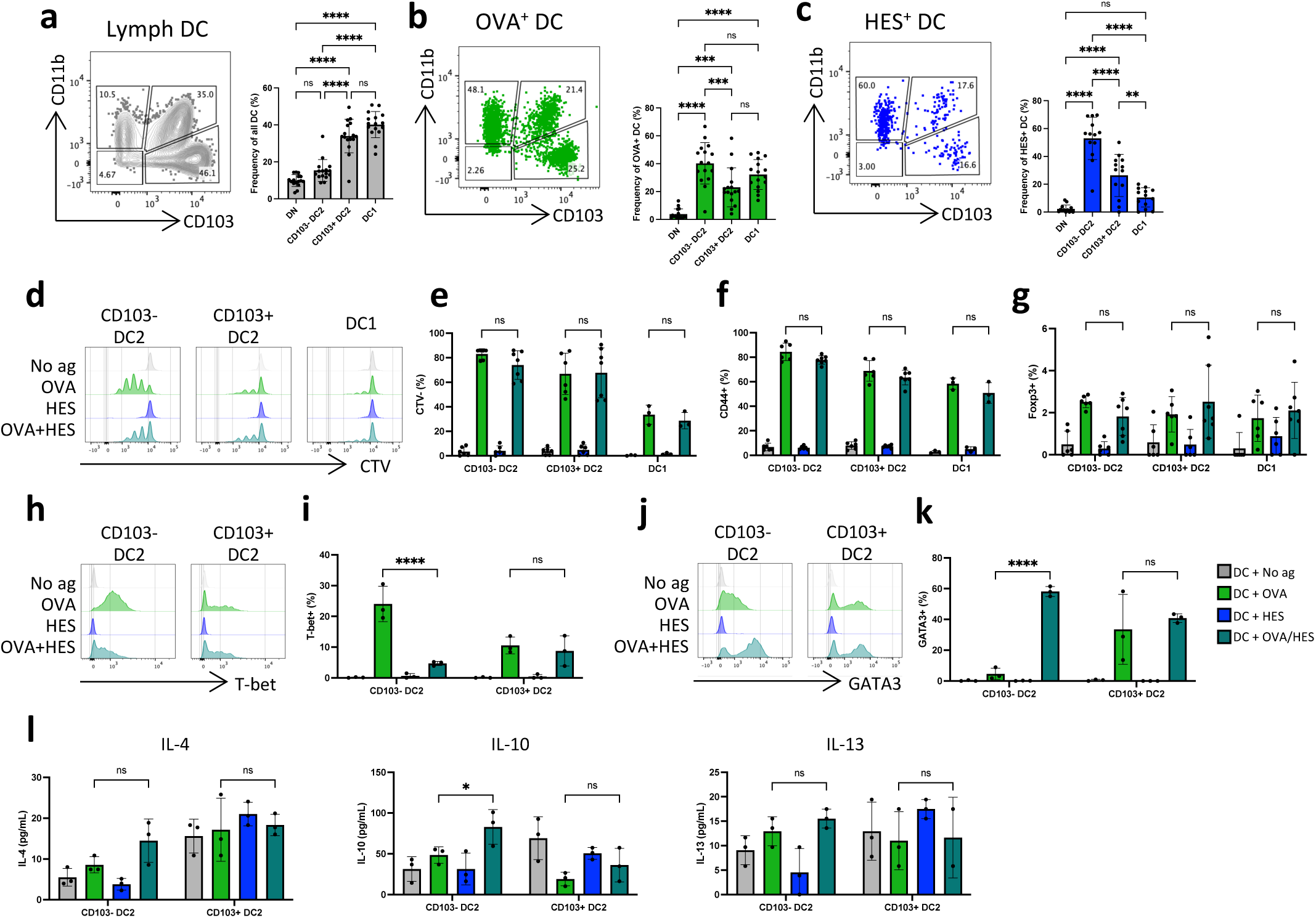
*Hpb* excretory-secretory products (HES) are trafficked, processed and presented differently between intestinal DC2 subsets. A) Representative contour plots (left) and proportional quantification (right) of DC subsets based on their CD103 and CD11b expression. Isolated from pseudo-afferent lymph by cannulation of the thoracic duct at steady state following a lymphadectomy of MLN. DCs were identified as live, single, CD45^+^, CD3^-^, B220^-^, CD64^-^, MHCII^+^ and CD11c^+^. B) Representative contour plots (left) and proportional quantification (right) of OVA^+^ DC subsets based on their CD103 and CD11b expression. Isolated from pseudo-afferent lymph by cannulation of the thoracic duct at steady state following a lymphadectomy of MLN. C) Representative contour plots (left) and proportional quantification (right) of HES^+^ DC subsets based on their CD103 and CD11b expression. Isolated from pseudo-afferent lymph by cannulation of the thoracic duct at steady state following a lymphadectomy of MLN. Data shown are pooled from 6 independent experiments (n=2-4 each). D) Representative histograms showing levels of cell trace violet (CTV) expression by naïve OTII CD4^+^ T cells cultured at a 10:1 ratio for 3 days with OVA and/or HES-pulsed DC subsets isolated SI LP at steady state. E) Quantified proportions of CTV^-^ within naïve OTII CD4^+^ T cells cultured at a 10:1 ratio for 3 days with OVA and/or HES-pulsed DC subsets isolated from SI LP at steady state. F) Quantified proportions of CD44^+^ within naïve OTII CD4^+^ T cells cultured at a 10:1 ratio for 3 days with OVA and/or HES-pulsed DC subsets isolated SI LP at steady state. F) Quantified proportions of Foxp3^+^ within naïve OTII CD4^+^ T cells cultured at a 10:1 ratio for 3 days with OVA and/or HES-pulsed DC subsets isolated from SI LP at steady state. Data shown are pooled from 2 independent experiments (n=3-4 each). H) Representative histograms of T-bet^+^ within naïve OTII CD4^+^ T cells cultured at a 10:1 ratio for 3 days with OVA and/or HES-pulsed DC subsets isolated SI LP at steady state. I) Quantification of T-bet^+^ within naïve OTII CD4^+^ T cells cultured at a 10:1 ratio for 3 days with OVA and/or HES-pulsed DC subsets isolated from SI LP at steady state. J) Representative histograms of GATA3^+^ within naïve OTII CD4^+^ T cells cultured at a 10:1 ratio for 3 days with OVA and/or HES-pulsed DC subsets isolated from SI LP at steady state. K) Quantification of T-bet^+^ within naïve OTII CD4^+^ T cells cultured at a 10:1 ratio for 3 days with OVA and/or HES-pulsed DC subsets isolated from SI LP at steady state. Data shown are representative of 2 independent experiments (n=3-4 each). L) Quantification of cytokines in the supernatant of naïve OTII CD4^+^ T cells cultured at a 10:1 ratio for 3 days with OVA and/or HES-pulsed DC subsets isolated SI LP at steady state. *p<0.05, **p<0.01, ***p<0.001,****p<0.0001 as assessed by two-way ANOVA with Šídák’s post-test correction for multiple comparisons. ns=not significant.

Having identified HES carrying potential amongst the DCs, we next tested the DC antigen presenting ability in context of HES antigens. In the absence of HES-specific TcR transgenic T cells to assess the antigen effect directly, we developed an antigen ‘piggyback’ system using OVA and OVA-specific T cell model that allowed the antigen presentation function of HES-loaded DC to be explored indirectly. We validated this system by quantifying how DCs take up fluorescently labelled antigens *in vitro* (Figure S2d). While DCs exposed to no antigen (grey dotplot and bar graph) served as a control, when DC were incubated with OVA, almost entirely all cells took it up (green dotplot and bar graph). When DCs were incubated with HES, the antigen was taken up at varying levels with around 60% having taken up some level of HES (blue dotplot and bar graph). Finally, when DC were incubated with both OVA and HES together, we identified that while approximately 60% had still taken up HES, all of the DC had also taken up OVA (aqua dotplot and bar graph). Taken together, we concluded that we can use this ‘antigen piggyback’ system with OVA specific T cells and possibly identify HES unique effects on the DCs, when comparing DCs incubated with only OVA or with HES and OVA. To do this, we sorted DC subsets from steady state SI LP, incubated them with OVA, HES or both OVA and HES and then co-cultured them with CTV-labelled, naïve, OVA specific CD4^+^ T cells.

First, we assessed T cell proliferation as measured by CTV dilution (Figure 3d). Under these conditions, we identified that while DCs with no antigen or incubated with only HES lacked an ability to induce T cell proliferation, both OVA and OVA+HES incubated DCs both were able to drive naïve T cells to proliferate at a similar rate (Figure 3e). Similarly, there were no changes observed in T cell activation as measured by CD44 expression 9Figure 3e) or induction of Tregs, measured by Foxp3 expression (Figure 3g). Next, we sought to determine the antigen loaded DC ability to polarise the differentiation of primed T cells by measuring T-bet and GATA3 expression, associated with Th1 and Th2, respectively (Figure 3h-k). Here, we identified that the addition of HES reduced the CD103^-^ DC2 ability to prime T cells to express T-bet, however the same effect was not observed in T cells co-cultured with CD103^+^ DC2 (Figure 3i). Conversely, CD103^-^ DC2 incubated with OVA+HES showed a significantly enhanced ability to simulate T cells to express GATA3, when compared to OVA-alone incubated CD103^-^ DC2 (Figure 3k). Like T-bet, regardless of HES, CD103^+^ DC2 ability to induce T cells to express GATA3 was not altered.

Finally, co-culture supernatant was also analysed for cytokines and while CD103^+^ DC2 did not change in their ability to stimulate T cell production of IL-4, IL-10, IL-13 regardless of what antigen they were incubated with; CD103^-^ DC2 incubated with HES in addition to OVA had an increased ability to induce T cell production of the cytokines (Figure 3l). Thus CD103^-^ DC2 population have a preferential ability to take up *Hp* derived HES antigens *in vivo* and exposure of CD103^-^ DC2 to HES *in vitro* increased their ability to skew T cells towards a Th2-like phenotype. In contrast, although CD103^+^ DC can also acquire HES *in vivo* and *in vitro*, exposure to HES did not change their T cell priming abilities. While these *in vitro* observations are line with the previously observed DC2 functional divergence during *Hpb* infection, it does not fully replicate the full functional profile of CD103^-^ and CD103^+^ DC2 *in vivo* (Figure 2), thus suggesting that the inflammatory environment during the infection may also play a role.

### DCs altered in proximity to the *Hp* granuloma

Given that antigen uptake alone could not explain the functional divergence between the CD103+ and CD103-DC2 populations observed during *Hpb* infection and evidence that some of the phenotypical changes correlate with known timepoint of *Hpb* lifecycle, we next hypothesised that worm might skew the intestinal environment in ways that result in the changes observed in the intestinal DC phenotype and function. A characteristic feature of *Hp* is that after invading from the intestinal lumen around 2DPI, the adult worm establishes itself within granuloma-like structures in the submucosa. These become most apparent around 7DPI, the time at which we found phenotypic and functional differences in DC. Therefore, we set out to characterise intestinal DC depending on their proximity to the Hp granuloma at 7DPI. To do this, we took advantage of the fact that at 7DPI, the granulomas approximately half of the size of the Peyer’s Patch and are visible by naked eye (Figure S3a). Thus, we dissected the individual granuloma and for comparison also isolated comparable size biopsies from the *Hpb* infected tissue without visible granuloma or naïve tissue. We isolated cells from the tissue and analysed the cellular composition. Previously, granuloma tissue has been described to contain large number of eosinophils, neutrophils and macrophages^33^ and markers associated with these cells were also confirmed to be enriched around the granuloma using microscopy (Figure S3b), thus we first set out to characterise myeloid cells in the tissue in order to validate our approach (Figure S4a). Analysis of cells isolated from granuloma-associated tissue showed a heterogeneous mixture of myeloid cells, comprising neutrophils, eosinophils, monocytes, macrophages and DCs. Proportionally, granuloma associated tissue showed an increase in neutrophils and monocytes (Figure S4b). Next, we quantified cell numbers and normalised it by weight of the tissue. In all cases, there was a higher abundance of myeloid cells in granuloma-associated tissue compared with *Hpb* infected non-granuloma tissue or non-infected SI LP. Thus, we concluded that this method allows for robust isolation of cells in the proximity of the granuloma.

Next, we used this approach to assess intestinal DC composition and explore if it is altered by the proximity to the granuloma (Figure 4a). As previously observed, we found that during *Hpb* infection the proportion of CD103^-^ DC2 is reduced, however this change was especially prominent in the tissue proximal to granuloma (Figure 4b). In parallel, the proportions of CD103^+^ DC2 were increased to the same extent in both infected tissues compared with control SI LP, while the proportion of DC1 was significantly increased only in the granuloma. Analysis of cell number per mg of tissue confirmed the significant increase in all DC subsets in *Hpb* SI LP we had observed previously, and this expansion was significantly greater for CD103^+^ DC2 and DC1 in the granuloma compared with non-granuloma SI LP (Figure 4c). Validation by imaging confirmed increased density of CD11c^+^CD11b^+^, CD11c^+^CD11b^+^ CD103^+^ and CD11c^+^CD103^+^ in direct proximity to the *Hpb* granuloma (Figure 4d).

**Figure 4.**
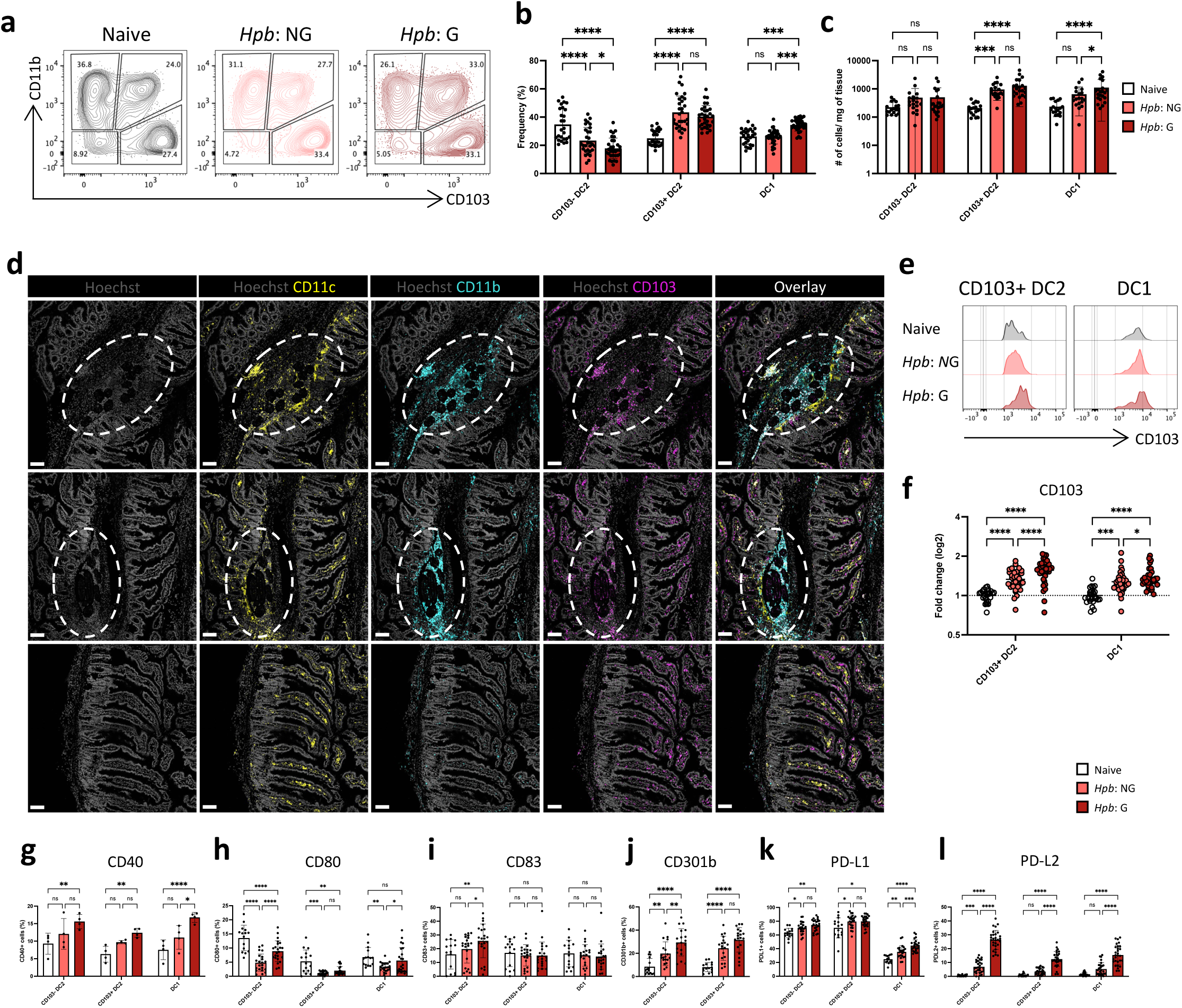
*Hpb* presence in the SI LP alters DC2 in subset specific manner. A) Representative contour plots of DC subsets based on their CD103 and CD11b expression. Isolated from SI LP at 7DPI with *Hpb* (either granuloma enriched tissue (G) or non-granuloma tissue (NG)) or naïve controls. DCs were identified as live, single, CD45^+^, CD3^-^, B220^-^, CD64^-^, MHCII^+^ and CD11c^+^. B) Quantified frequency of DC subsets isolated from SI LP at 7DPI with *Hpb* (either granuloma enriched tissue (G) or non-granuloma tissue (NG)) or naïve controls. Data shown are pooled from 6 independent experiments (n=2-6 each) C) Quantified absolute cell number of DC subsets isolated from SI LP at 7DPI with *Hpb* (either granuloma enriched tissue (G) or non-granuloma tissue (NG)) or naïve controls. Data shown are pooled from 6 independent experiments (n=2-6 each). D) Imaging of granuloma (top, middle panels) and non-granuloma (bottom panel) tissue stained for nuclear (Hoechst), CD11c, CD11b and CD103. Shown as individual stains and then all overlayed. E) Representative histogram of CD103 expression in DC subsets isolated from SI LP at 7DPI with *Hpb* (either granuloma enriched tissue (G) or non-granuloma tissue (NG)) or naïve controls. F) Quantification of fold change of CD103 expression (MFI) in DC subsets isolated from SI LP at 7DPI with *Hpb* (either granuloma enriched tissue (G) or non-granuloma tissue (NG)) or naïve controls. Data shown are pooled from 6 independent experiments (n=4-6 each) G) Quantification of CD40^+^(G), CD80^+^(H), CD83^+^(I), CD301b^+^(J), PD-L1^+^(K), PD-L2^+^(L) within DC subsets isolated from SI LP at 7DPI with *Hpb* (either granuloma enriched tissue (G) or non-granuloma tissue (NG)) or naïve controls. Data shown are pooled from 1-4 independent experiments (n=4-6 each).

Importantly, changes in expression of several markers previously identified (Figure 1) appeared to be location specific (Figure 4e-l). In particular, the previously observed increase in the levels of CD103 expression on CD103^+^ DC2 was particularly striking in the cells isolated from the granuloma-associated tissue (Figure 4e, f). A similar, but less strong change was also observed in DC1. Next, we assessed expression patterns of CD40, CD80, CD83, CD301b, PD-L1 and PD-L2 as these were previously seen to change total *Hpb* SI LP (Figure 4g-l). Here, we observed that the proportion of cells positive for the markers was significantly greater in all DC populations isolated from granuloma compared with non-granulomatous SI LP. The proportion of CD40^+^ CD103^-^ DC2 was specifically increased in cells from granuloma-associated tissue (Figure 4g). Interestingly, the proportion of CD40^+^ CD103^+^ DC2 previously was not observed to change, however fractioning based on location revealed a small but significant increase in cells proximal to granuloma. Furthermore, the decreased proportion of CD80^+^ within the CD103^-^ DC2 population seen in total SI LP was partly, but significantly, restored when these subsets were examined in context to the proximity of granuloma (Figure 4h). This was not seen with CD103+ DC2, where the proportions expressing CD80 were significantly reduced compared with control SI LP, regardless of their proximity to the worm. Similarly, proportional increase of CD83^+^ CD103^-^ DC2 was restricted to the cells in the granuloma (Figure 4i), and the proportional increase of CD301b^+^, PD-L1, and PD-L2 was granuloma-associated (Figure 4j, k,l). In case of CD301b and PD-L2, this increase was particularly significant within the CD103^-^ DC2 population.

Together these results reveal that previously observed changes in DC phenotype can be associated with their location, with cells in close proximity to the worm showing a distinct phenotypic signature.

### The transcriptional profile of DC2 subsets in Hp infection is dependent on their proximity to the worm

Having identified phenotypically and functionally distinct populations of DC during *Hpb* infection that were specifically located in close proximity to the granuloma, we sought to identify genes and signalling pathways underlying this functional divergence in the DC2 populations. To this end, we performed bulk RNA sequencing of FACS-purified CD103- and CD103+ DC2 populations isolated from naïve SI LP as well as granuloma and non-granuloma regions at 7DPI with *Hpb* (Figure 5).

**Figure 5.**
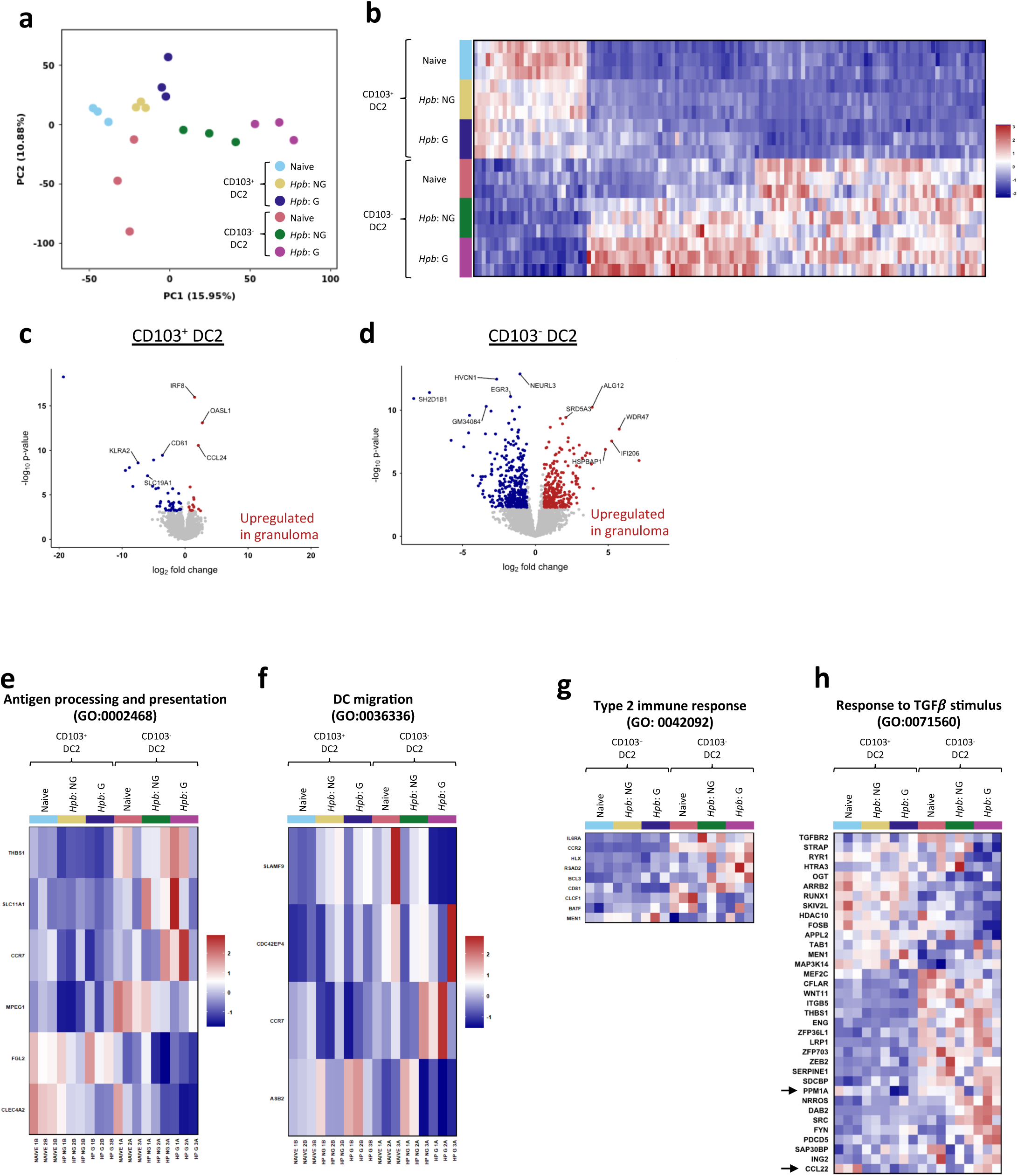
Transcriptional profile of DC2 subsets based on their proximity to granuloma at 7 DPI with *Hpb*. A). PCA plot of DC subsets based on their CD103 and CD11b expression. Isolated from SI LP at 7DPI with *Hpb* (either granuloma enriched tissue (G) or non-granuloma tissue (NG)) or naïve controls. DCs were identified as live, single, CD45^+^, CD3^-^, B220^-^, CD64^-^, MHCII^+^ and CD11c^+^. B) Heatmap of DEGs within DC subsets based on their CD103 and CD11b expression. Isolated from SI LP at 7DPI with *Hpb* (either granuloma enriched tissue (G) or non-granuloma tissue (NG)) or naïve controls. C) Pairwise comparison of DEGs between CD103^+^ DC2 isolated from SI LP at 7DPI with *Hpb* (either granuloma enriched tissue (G) or non-granuloma tissue (NG)). D) Pairwise comparison of DEGs between CD103^-^ DC2 isolated from SI LP at 7DPI with *Hpb* (either granuloma enriched tissue (G) or non-granuloma tissue (NG)). E-H) DEGs that reach significance in GO terms 0002468, 0036336, 0042092, 0071560.

Initial principal component analysis indicated clear clustering of groups based on DC2 population, presence of infection and location (Figure 5a). There was a large amount of differentially expressed genes (DEGs), particularly in the CD103^-^ DC2 population isolated from granuloma-associated tissue (Figure 5b). To identify genes uniquely associated with granuloma residence we performed pair-wise comparisons of the transcriptomes of the individual DC2 populations isolated from granulomatous or non-granulomatous regions of infected intestine (Figure 5c,d). This showed that of the relatively few DEGs for granuloma-associated CD103^+^ DC2 subset, the most significantly upregulated were *Irf8*, involved in DC development, *Oasl1*, an interferon-inducible gene^34^ and chemokine *Ccl24*^435^, implicated in allergic responses. The larger group of downregulated DEGs included those encoding the *Klra2*, shown to be upregulated in myeloid cells in response to various inflammatory stimuli^36^, *Slc19A1*, cGAMP transporter upstream of STING signalling pathway^37^ and the tetraspanin *Cd81* implicated in migration, antigen processing and presentation^38^. The pattern of DEGs in granuloma-associated CD103-DC2 subset was very different. Amongst those upregulated included genes encoding the mannosyl-transferase enzyme *Alg12* and steroid 5-alpha reductase 3 enzyme *Srd5A3* involved in glycosylation^39,40^, interferon gamma inducible protein *Ifi206, Wdr47* implied in cytoskeleton remodelling^41^, while decrease was observed in genes such as *Egr3* implicated in cell anergy^42^ and *Hvcn1* shown to decrease in response to DC activation^43^.

As this unbiased identification of DEGs did not immediately identify clear immune pathways associated with DC2 located in granulomata, we decided to employ more targeted approach using previously defined immune signatures (Figure 5e-h). First, we screened our data for gene signatures associated with antigen processing and presentation (GO:0002468) or DC maturation and migration (GO:0036336) (Figure 5e, f). Within the antigen processing signature genes, CD103^-^ DC2 isolated from granuloma-associated tissue were enriched for the *Thbs1* gene encoding thrombospondin, the cation transporter *Slc11a1* that plays a role in antigen presentation and DC maturation^44^ and the chemokine receptor *Ccr7*^45^. Genes associated with this pathway that were decreased in granuloma-associated CD103^-^ DC2 included *Mpeg1* that has been implicated to be necessary for antigen presentation to CD4+ T cells^46^. In parallel, CD103^+^ DC2 isolated from granuloma-associated tissue were enriched for *Fgl2,* which has been implicated to have immunomodulatory properties and inhibitory effect on DC maturation^47^ and had decreased expression of *Clec4a2* encoding DCIR implicated in regulation of DC proliferation^48^ (Figure 4e). Within the DC migration signature, CD103^-^ DC2 from the granuloma-associated tissue showed a decrease in expression of *Slamf9* previously implicated to negatively regulate myeloid cell migration^49^. Interestingly, the *Asb2* gene that has been shown to be highly expressed in immature DC and required for their migration, while downregulated upon DC maturation^50^ appeared to be increased in granuloma-associated CD103^+^ DC2, but decreased in the CD103^-^ DC2, suggesting possible divergence between the two DC2 populations. Next, we screened our data for transcriptional changes associated with Type 2 immune responses (GO: 0042092) (Figure 5g). Genes associated with this signature were mostly enriched in the CD103^-^ DC2 isolated from the proximity of granuloma.

Together, these data show that CD103^+^ and CD103^-^ DC2 are transcriptionally distinct and appear to be in different activation/maturation states based on the proximity to the granuloma. Here, in line with our previous observations in phenotype and function, CD103-DC2 appear to be more transcriptionally activated, mature and involved in driving of pro-inflammatory immune response. However, CD103^+^ DC2 transcriptional profile is ambiguous with less clear association with immune function.

Next, we set out to identify a possible source of this transcriptional and functional divergence in the intestinal DC2 during *Hpb* infection. TGFβ signalling is the most widely described immunomodulatory route for *Hpb*^51–55^. Furthermore, TGFβ signalling has also been previously described to be crucial for CD103^+^ DC2 development at homeostasis^56^. Thus, we first set out to screen our dataset for transcriptional changes associated with response to TGFβ stimulus (GO:0071560). Most notably, some changes were DC2-population and spatial-origin specific, as in case of *Ppm1a*, a SMAD2 phosphatase that negatively regulates TGFβ signalling^57^, and *Ccl22*, a chemokine associated with driving type 2 immunity^58,59^, that decreased in granuloma-proximal CD103^+^ DC, yet increased in the CD103^-^ DC2 population (Figure 5h).

Together, the transcriptomic profiling of DC2 populations in different locations in context of *Hp* granulomas showed that CD103^-^ DC2 are predominantly activated, meantime CD103^+^ DC2 to have immature or regulatory transcriptional profile. Furthermore, both populations showed changes that can be associated with TGFβ stimulus, however they were divergent between the populations possibly explaining the functional and phenotypical divergence previously observed.

### TGF*β*-signalling skew DC development towards immature and regulatory phenotype

Having identified the distinct differences in TGF*β* signalling lead us to hypothesise that the heterogeneous DC2 populations found during *Hp* infection might be a result of skewed precursor development, possibly by endogenous TGF*β* signalling or TGF*β* mimic (TGM) molecules produced by the helminth. To explore this idea, we cultured bone marrow derived, FACS-purified DC precursors in presence of TGF*β* or purified TGM. First, we assessed DC phenotype based on changes we had observed during *in vivo* infection model (Figure 6a). We identified that pre-DCs derived in presence of TGF*β* or TGM had a lower levels of co-stimulatory molecules CD80 and CD86, reminiscent of CD103^+^ DC2 phenotype observed *in vivo* (Figure 1e). Additionally, these cells also had lower levels of MHCII and IRF4 associated with more immature state. Finally, both, the TGF*β* and TGM, treatments increased IRF8 levels in the developing pre-DCs *in vitro*, similar to the significant increase in the granuloma associated CD103^+^ DC2 *in vivo* seen in the transcriptional analysis (Figure 5c). Next, we assessed the T cell stimulating abilities of the TGF*β* modulated pre-DCs by co-culturing them with naïve T cells. Here, we identified that DCs derived in presence of TGF*β* had a reduced ability to stimulate T cell proliferation, as measured by CTV (Figure 6b). This observation is in line with the more immature phenotype of the *in vitro* derived pre-DC. However, we observed that these cells had an increased ability to drive naïve T cells to develop into Foxp3^+^. Thus, we concluded that the regulatory, immature phenotype of the CD103^+^ DC2 found *in vivo* during Hp infection might be result of TGF*β* signalling and likely a result of direct immunomodulatory effect of the TGF*β* mimic molecules secreted by *Hp*.

**Figure 6.**
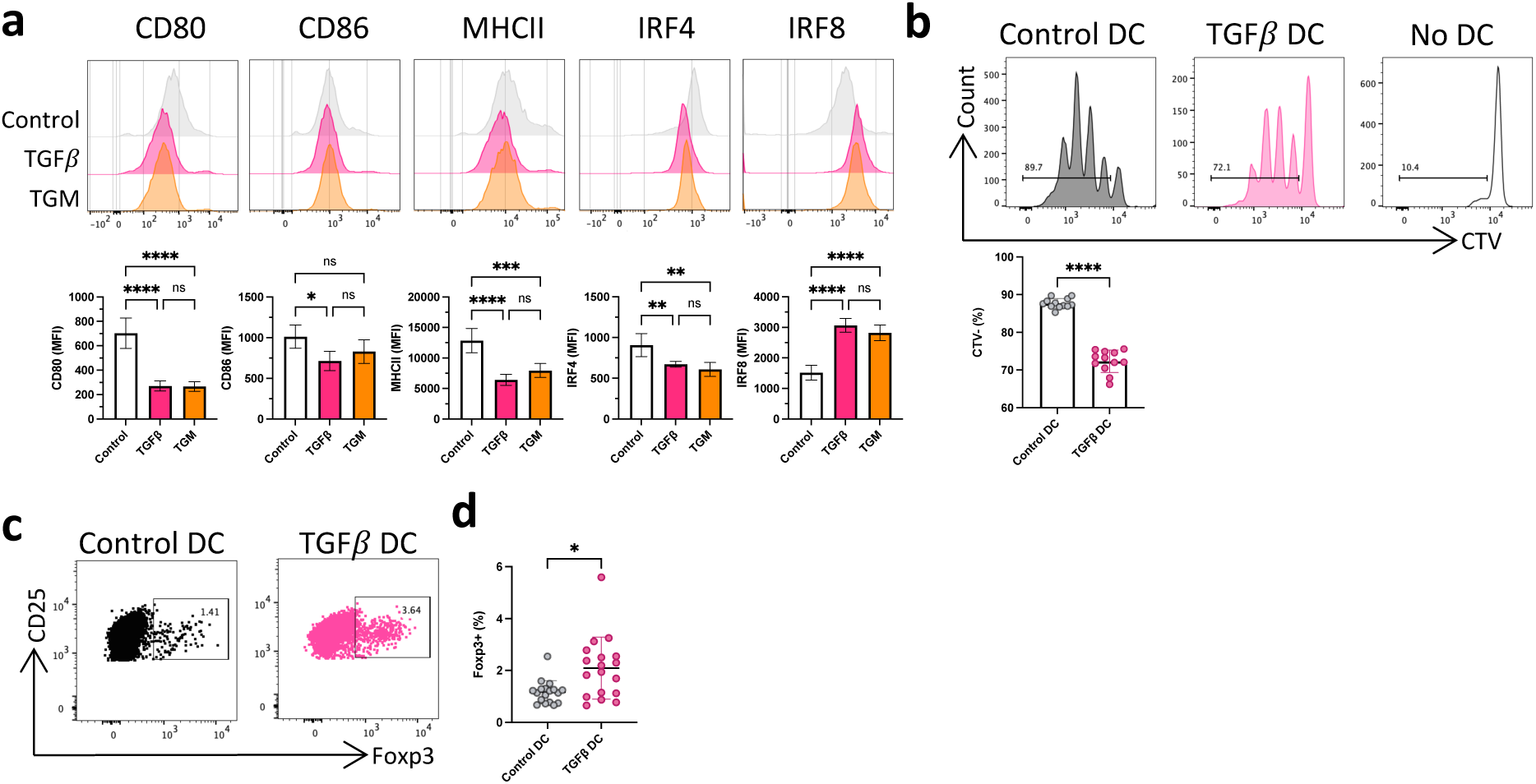
TGF*β* signalling skew DC development towards immature and regulatory phenotype *in vitro*. A) Representative histograms of CD80, CD86, MHCII, IRF and IRF8 expression in FACS sorted pre-cDC that were cultured for 4 days in complete medium containing 400ng/mL Flt3 ligand and 20ng/mL GM-CSF(Control) and 200ng/mL TGFβ or 200ng/mL TGM. Gated on live CD45^+^CD64^-^Ly6C^-^CD11c^+^MHCII^+^ cells. Data shown are from 1 experiment (n=4-5 each). B) Representative histogram (top panel) and quantification (bottom panel) of CTV^-^ within naïve OTII CD4^+^ T cells cultured at a 10:1 ratio for 3 days with OVA-pulsed pre-cDC in presence of TGFβ. Representative dotplot (C) and quantification (D) of Foxp3^+^ within naïve OTII CD4^+^ T cells cultured at a 10:1 ratio for 3 days with OVA-pulsed pre-cDC in presence of TGFβ. *p<0.05, ***p<0.001, ****p<0.0001, ns =not significant, as assessed by one-way ANOVA with Tukey’s post-test for multiple comparisons (A) or two-tailed, unpaired Student’s *t*-test (B, D).

## Discussion

Here, we have presented the first-ever in-depth characterisation of intestinal DCs during intestinal helminth *Hpb* infection. As DC2 has been previously described to drive Th2 responses in other helminth infection models^8,12,32^, we focused our analysis on this subset of intestinal DCs. Our work reveals an emergence of complex SI LP DC2 compartment that dynamically changes in abundance, phenotype and function over the course of *Hpb* infection. In particular, anti-helminth immune response appears to be primary driven by CD103^-^ DC2 population. This populations adapted a phenotype previously associated with Th2-driving abilities, such as increased expression of CD40, CD301b, IRF4, PD-L2 (Figure 1d). At 7DPI *ex vivo* these cells drove T cell activation and increased production of cytokines IL-13 and IL-10, that can be associated with Th2^7^ (Figure 2). (ref ref). Interestingly, the phenotypical change was especially striking in the cells isolated from the proximity of *Hpb* granuloma. Finally, these same molecules as well as other co-stimulatory markers, such as CD80 and CD83, were particularly increased on CD103-DC2 isolated from the proximity of *Hpb* granuloma (Figure 4). Furthermore, throughout the course of infection, these molecules were upregulated to a much higher level when compared to the CD103^+^ DC2 counterparts (Figure S1d-h), especially between the timepoints when *Hpb* is present in the gut wall in order to complete its maturation (Figure S1d). CD103^+^ DC2 not only had a less severe change in the phenotype (Figure 1e), but they also failed to drive strong effector response and instead, remained to be superior drivers of Tregs (Figure 2).

Furthermore, in steady state CD103^-^ DC2 were better at carrying and processing *Hpb* antigens (Figure 3) and the transcriptional analysis of DC2 populations showed that CD103^-^ DC2 in the near proximity to the granuloma had gene signatures associated with glycosylation, which could hint towards processing of *Hpb* antigens^60^, that have been previously described to be glycoproteins^30^.

As the divergence within the two distinct CD103^-^ and CD103^+^ DC2 populations was associated with the proximity to the *Hpb* granuloma (Figure 4), we hypothesised that *Hpb* itself or the micro-environment present during infection could be the source of the signals that determine the DC2 phenotypical and functional heterogeneity. Transcriptional analysis confirmed this and identified TGF*β* signalling as one of the possible sources of the SI LP DC2 population divergence during *Hpb* infection (Figure 5). Subsequent *in vitro* work confirmed a role for TGF*β* signalling in skewing the DC development (Figure 6). Supplementation of TGF*β* or worm-produced TGM partly replicated phenotype and function of CD103^+^ DC2 found *in vivo* during *Hpb* infection. Together, this data provides a mechanistical explanation of how *Hpb* parasite hijack the endogenous TGF*β* signalling to inhibit or skew DC2 development into more immature, regulatory cells. This immunomodulation of SI LP DC2 results in overall weaker initial anti-helminth Th2 response thus providing a route of immune evasion and the necessary protection to the helminth necessary for its maturation and eventual exit from the gut tissue.

CD103^+^ DC ability to drive Tregs via TGF*β* and RA dependant mechanism is well described^29^ and thus not novel, however a pathogens ability to hijack these endogenous DC signalling pathways as a route of immunomodulation has never been described before. Thus, our work builds on the growing evidence of helminth ability to immunomodulate various aspect of host immune system in order to prevent expulsion^5^. While others have showed *Hpb* effect on Tregs, our work provide a much more nuanced mechanistic explanation that encompasses an effect on the whole adaptive side of the immune response.

Furthermore, in addition to novelty regarding helminth infection biology, our findings are also informative in much larger context of routes of immune evasion. Understanding how the microenvironment can shape DC phenotype, development and function has been of increasing interest in the field of tumour biology. There has been a particular focus on TGF*β* signalling in the tumour micro environment (TME) with it being implicated as a source of immune evasion in tumours associated with immune suppression, especially in context of DCs^61–63^. Thus, the similar findings in our model suggest an overarching theme of basic biology bridging these two fields.

## Methods

### Animals

C57BL/6 females were purchased from Envigo at 5-6 weeks of age and used for procedures at 7-8 weeks of age. Mice were housed under specific pathogen free conditions at the Common Research Facility, University of Glasgow. OTII CD45.1 mice were a gift from Dr Ed Roberts (Beatson Cancer Research Institute, Glasgow). Age matched animals were used for all experiments and all procedures were carried out under personal and project licences issued by the UK Home Office.

### Infection with *Hpb*

Mice were infected with 200 *Hpb* L3 larvae (a gift from Professor Rick Maizels) by oral gavage and tissues were harvested 2-14 days post infection.

### Isolation of lamina propria cells

Small intestine (SI) were removed, cut open longitudinally and rinsed in cold PBS, with Peyer’s patches being removed from the small intestines. Intestinal tissue was enzymatically digested as described previously^6^. In brief, the epithelium and mucus were first removed by washing tissues 3 times in warm calcium/magnesium free HBSS containing 2mM EDTA (Gibco) for 15 minutes at 37°C with shaking at 250rpm. Small intestines were then digested by incubation with 0.5mg/mL collagenase VIII (Sigma), while colons were digested with 0.5mg/mL collagenase V (Sigma), 0.65mg/mL collagenase D (Roche), 1mg/mL Dispase (Gibco) and 30ug/mL DNase (Roche) each for 15-20 minutes at 37°C with shaking. Digestion was stopped by adding cold complete RPMI medium (RPMI 1640 contained 10% heat inactivated foetal calf serum (FCS), 2mM L-glutamine, 100μm/mL penicillin/streptomycin and 50μm b-mercaptoethanol (all Gibco). Cells were then passed through a 100μm and then a 40μm mesh EASYstrainer (Greiner), before being washed twice with cold complete RPMI medium for 10 minutes in 4°C at 400g at stored at 4°C before use. For separation of individual granuloma-associated tissue, SI was processed the same way, but visible granuloma tissue were removed using dissection scissors. For each granuloma, a comparable size non-granuloma tissue in the proximity to the paired granuloma was removed for control. Comparable size naïve tissue was removed at the same location down the length of the SI from naïve animals to be used as control. Tissue was processed, washed and enzymatically digested in the same manner as the whole gut tissue. Each individual collection tube was weighed before and after tissue collection to determine the tissue weight that was used for normalisation.

### Isolation of lamina propria cells

Small intestine draining mesenteric lymph nodes (sMLN) and spleens were cut into small pieces and placed in enzymatic digest medium containing 1mg/mL collagenase D (Roche) for 40 minutes at 37°C with shaking at 250rpm. Digestion was stopped by adding cold complete RPMI. Cells were then passed through a 100μm and then a 40μm mesh EASYstrainer (Greiner), before being washed twice with cold complete RPMI medium for 10 minutes in 4°C at 400g at stored at 4°C before use.

### Generation of cDC from bone marrow precursors *in vitro*

Bone marrow cells were isolated from femurs and tibiae of 3-4 adult mice per experiment. Red blood cells were lysed using ACK lysing buffer (Thermo Fisher) for 1 minute and cells were washed with FACS buffer for 10 minutes at 4°C at 400g. Pre-cDC were then purified by first enriching using a MagniSort^TM^ Magnet (eBioscience) and MagniSort^TM^ SAV Negative Selection Beads (eBioscience) with biotinylated anti-CD3, anti-CD19, anti-Ly6G and anti-B220 at a 1:300 dilution. The resulting cells were then FACS sorted on an BD FACSAria^TM^ III using anti-CD3-BV605(1:100), anti-CD19-BV605 (1:100), anti-NK1.1-PercP-Cy5.5 (1:100) and anti-CD135-biotin in combination with PE-streptavadin (1:200) (all from eBioscience) and anti-CCR9-PE-Cy7 (1:100), anti-B220 (1:200), anti-CD11c-FITC(1:100), anti-CD11b-APC-Cy7 (1:200) and anti-MHCII-BV421 (1:200) (all BioLegend).

Aliquots of approximately 15,000 pre-cDCs were then cultured for 4 days in a total volume of 150μl in flat-bottomed 96-well plates (Costar, #3596) containing complete RPMI medium supplemented with 20ng/mL GM-CSF (Peprotech) and 400ng/mL Flt3L (Peprotech), together with 200ng/ml TGFβ (Prepotech) or TGFβ mimic (TGM) (purified in Rick Maizels laboratory). Cells were cultured at 37°C and 5% CO2.

### Flow cytometry

Aliquots of 5-10 x 106 cells were first assessed for viability staining using eFlour780 (eBioscience) fixable viability dye (1:1000) or 7AAD (Biolegend) viability dye (1:100). Non-specific binding to FC receptors was prevented by staining with anti-mouse CD16/32 (eBioscience) (1:100) for 20 mins at 4°C. Cells were then stained using the FACS antibodies at a 1:200 dilution in FACS buffer (DPBS supplemented with 2mM EDTA (Gibco) and 2% FBS(Sigma)) unless specified otherwise (Table 1) for 30 mins at 4°C. For intracellular staining of nuclear transcription factors and histone modifications, the eBioscience Foxp3 Transcription Factor Staining Buffer Set was used according to the manufacturer’s instructions. In brief, cells were fixed using 200μL fixation solution for 1 hour at room temperature before being washed with permeabilization buffer for 5 minutes in 20°C at 400g. Intracellular staining was performed for 1 hr at room temperature. Cells were analysed on BD Fortessa and FlowJo software (BD). Expression of proteins was assessed either as a % positive cells of the parent population or as median fluorescence intensity (MFI). ALDEFLOUR (# 01700, STEMCELL Technologies) assay was performed according to manufacturer’s instructions.

**Table 1.**
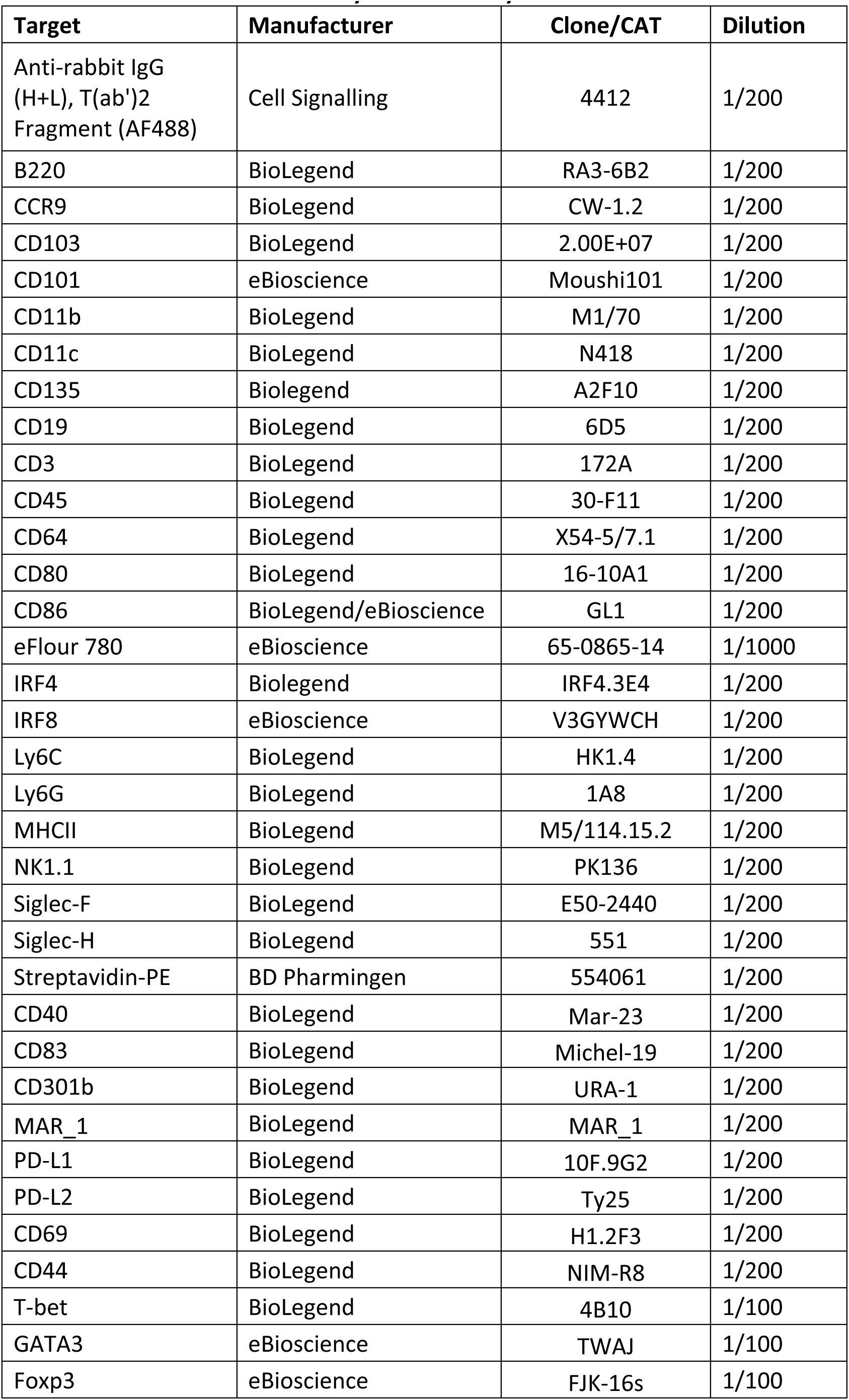
Antibodies used in flow cytometric analysis.

### Co-culture of DC and T cells

Naïve, OVA-specific CD4^+^ T cells were purified from the spleen and lymph nodes of OTII TCR transgenic mice using the MojoSort^TM^ Mouse CD4 Naïve T cell isolation Kit according to the manufacturer’s instructions (Biolegend). Cells were then stained with the CellTrace^TM^ Violet Cell proliferation kit (ThermoFisher) according to the manufacturer’s instructions. FACS-isolated colonic cDC populations or *in vitro* generated DC were pulsed with OVA protein (#SLBS4311, Sigma) at 2mg/mL, HES (purified in Rick Maizels laboratory) at xxx concentration or both for 2 hours at 37°C and co-cultured with T cells at 1:10 ratio for 3 days in complete RPMI in a total volume of 300μl in round-bottomed 96-well plates (Costar, #3799).

### Cytokine analysis

Supernatant of co-cultures was stored at -20C and analysed using DuoSet ELISA kits (Bio-Techne, #DY413 and #DY404) for IL-4 and IL-13 and BD™ Cytometric Bead Array (CBA) Mouse Inflammation Kit (BD, #552364) for IL-2, IL-6, IL-10, IL-17A, INFγ and TNFα according to manufacturers’ instructions.

### Antigen uptake and DC migration

Isolation of DC within the pseudo-afferent lymph was performed as previously described^31,64^. OVA (#SLBS4311, Sigma) or HES (purified in Rick Maizels laboratory) were fluorescently conjugated using Alexa Fluor™ 647 or 488 Microscale Protein Labeling Kit (A30009 and A30006 , ThermoFisher) according to the manufacturers instructions. At the laparotomy necessary for thoracic duct cannulation, up to 25μl fluorescently conjugated OVA or HES were injected within the SI LP. After 16 hrs, lymph was collected, washed x2, red blood cells were lysed using ACK lysing buffer (Thermo Fisher) for 1 minute and cells were subsequently processed for flow cytometry analysis.

### Bulk RNA sequencing

Cells isolate from granuloma, non-granuloma and naïve SI LP tissue were FACS-isoalted on an BD FACSAria^TM^ III using anti-CD3(1:100), anti-CD19-BV605 (1:100), anti-NK1.1 (1:100), anti-B220 (1:200), anti-CD64 (1:200), and anti-MHCII (1:200), anti-CD11c (1:100), anti-CD11b (1:200) and anti-CD103 (1:200). Each sample was pooled from 4 individual animals. Immediately after cell sort, cells were washed and resuspended in RLT supplemented with 0.1% β-ME. Cells were lysed by vigorous vortex for 1 min and flash-frozen using dry ice. RNA was extracted immediately after using RNeasy Plus Micro Kit (Qiagen) according to the manufacturer’s instructions. RNA library preparation and sequencing was performed commercially by Novogene and sequenced using NovaSeq X Plus (PE150) platform.

### Bioinformatic analysis

RNA-Seq fastQ files were quality control checked using FastQC (https://www.bioinformatics.babraham.ac.uk/projects/fastqc/, v0.11.7) and aligned to the Mus musculus reference genome (GRCm39, vM30) using STAR^65^ (v 2.7.9a). Genes with a mean expression <1 read per sample were removed from merged counts files. Expression and differential expression values were generated using DESeq2^66^ (v 1.42.1) using a pairwise model with no additional covariates. Genes with a median expression < 20 across samples were excluded from downstream analysis. The processed data was then visualised using RStudio and Searchlicht2^67^ according to the multiple differential expression (MDE) workflow using an absolute log2-fold cut-off of 0.58 and an adjusted p-value of 0.05 across all comparisons. Gene ontology (GO) annotation pathways of interest (GO:0002468, GO:0036336, GO: 0042092, GO:0071560; https://www.informatics.jax.org/vocab/gene_ontology/) used to generate heatmaps for MDE comparisons.

### Multiplex immunofluorescent imaging

Fresh Frozen OTC-embedded gut rolls of the upper half of the small intestine sections were prepared as previously described^68^, in short - small intestine samples were harvested in PBS, then snap frozen in TissueTek™ OCT™ (Sakura), block were stored at -80C. 10um sections were mounted on coverslips precoated in Poly-L-lysine (Sigma).For CODEX staining and imaging, oligonucleotide sequences were purchased from Integrated DNA Technologies and conjugated to primary antibodies as previously described ^69,70^(Table 2). Coverslip-mounted samples were defrosted at room temperature for 2mins, then incubated in acetone for 10min then rehydrated for 10min in SME (0.5% BSA, 5mM EDTA in PBS) following 2min of airdrying. Samples were then fixed in 1.6% formaldehyde in SME, washed twice with SME, incubated in EQ buffer (50% v/v SME, 0.25M NaCl, 0.1M Phosphate buffer in dH_2_O) for 10mins, then incubated at room temperature for 30min in blocking buffer (50ug/ml mouse IgG (Sigma), 50ug/ml rat IgG (Sigma), 0.5mg/ml sheared salmon sperm DNA (ThermoFisher), 3.5uM of each blocking oligonucleotide in EQ buffer). Conjugated antibodies diluted in blocking buffer were then added and left overnight at 4°C with gentle rocking. Following three washes in EQB buffer, antibodies were fixed for 10 min in 1.6% formaldehyde in SME05 (0.5M NaCl in SME), followed by 2 washes in PBS, then incubated for 5min in ice-cold methanol. This was washed twice in PBS, then samples were incubated in 3mg/mL bis(sulfosuccinimidyl)suberate crosslinker (BS3, ThermoFisher), washed twice in PBS, then stored in in SME05 at 4°C until imaging. Stitched images of Z-stacks were obtained using a Nikon A1R confocal microscope. Coverslips were sealed on a customised stage attached to a buffer exchange device to allow multicycle imaging, which involves cycles of applying the secondary oligo mix, imaging, stripping the signal with 80% DMSO, then washing twice with 20% DMSO to repeat the cycle for the next secondary oligo mix.

**Table 2.**
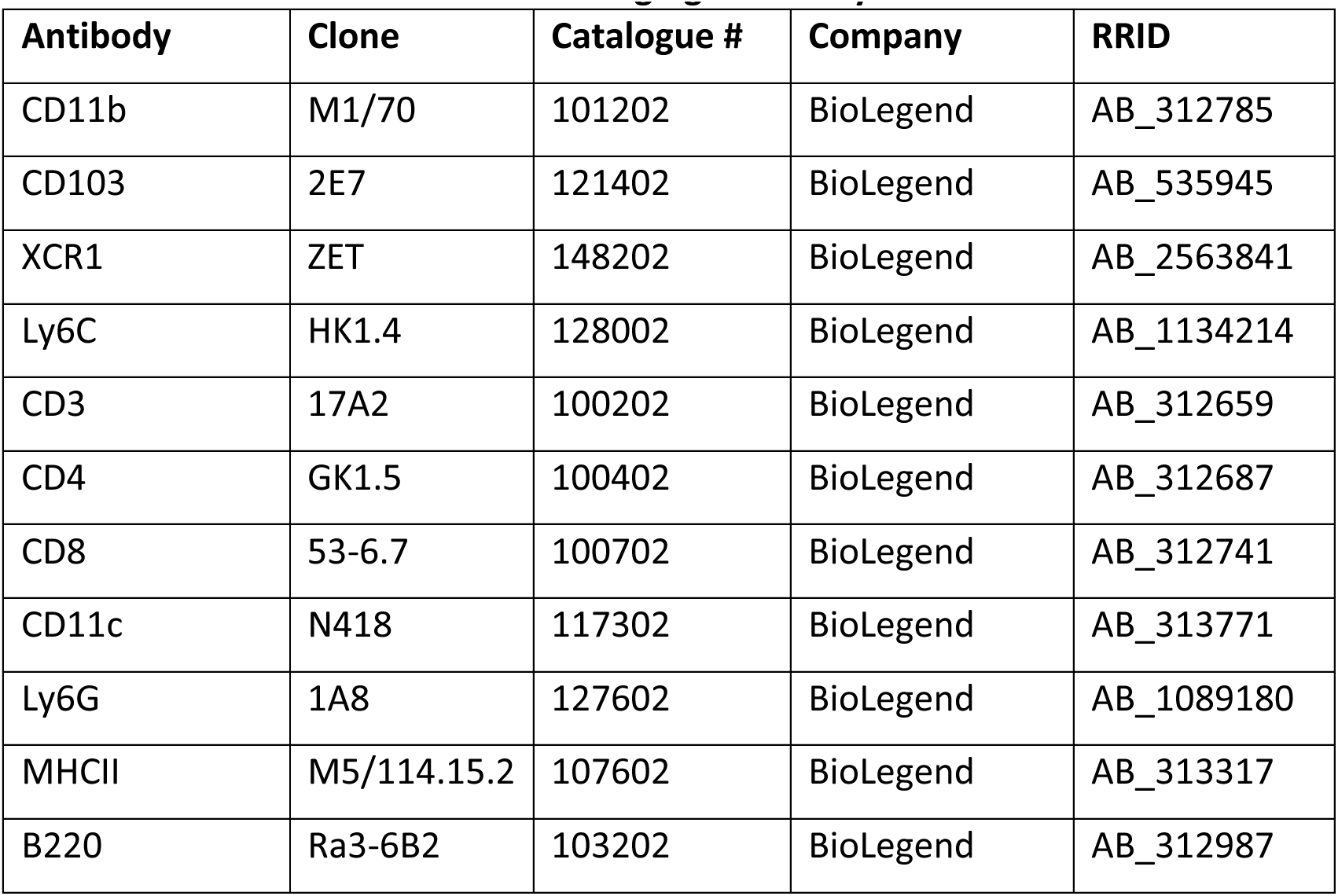
Antibodies used in CODEX imaging and analysis.

### Statistical analysis

Prism (GraphPad) software was used for all statistical analysis. Data were tested for normal distribution using the Shapiro-Wilk test for normality. Statistical differences were calculated using either one-way ANOVA (with Dunnett’s or Tukey’s post-test for multiple comparisons), two-way ANOVA (with Šídák’s post-test for multiple comparison) or Student’s *t* test, with the data being plotted as means and SD. Results were considered significant based on their *p*-values (*p<0.05, **p<0.01, ***p<0.001, ****p<0.0001).

### 2.12 Illustrations

Summary illustration was created using BioRender.com with an academic subscription. Publication license number PO25GQABJP

## Notes

### Competing Interest Statement

The authors have declared no competing interest.

