## Supplementary material for "Intestinal helminth skews DC2 development towards regulatory phenotype to counter the anti-helminth immune response"

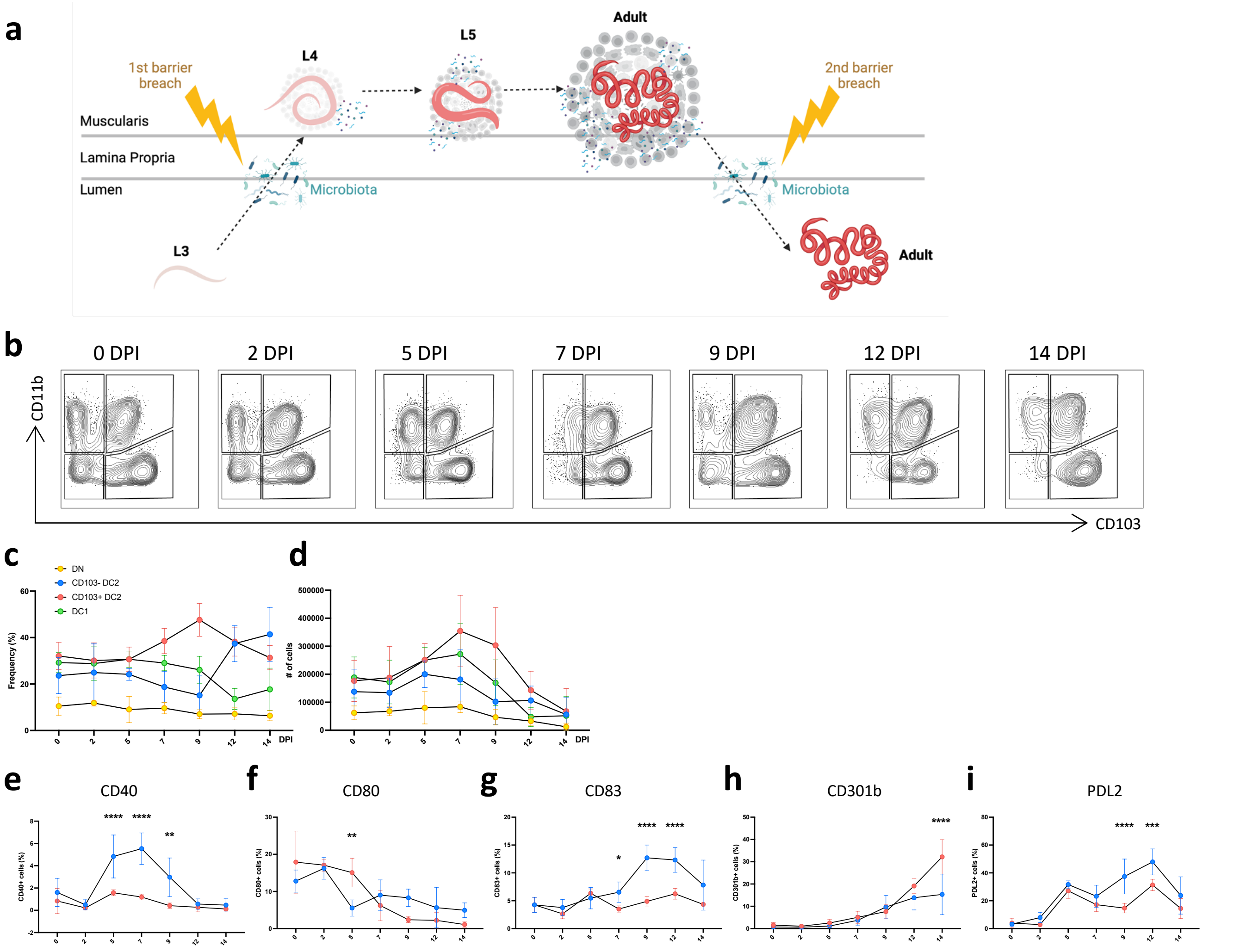


**Figure S1. Small intestinal lamina propria (SI LP) DCs over the course of *Hpb* infection (0-14DPI).**
A) Illustration of *Hpb* lifecycle 0-14 DPI. B) Representative contour plots of DC subsets based on their CD103 and CD11b expression. Isolated from SI LP at 0-14 DPI with *Hpb* or naïve controls. DCs were identified as live, single, CD45$^+$, CD3$^-$, B220$^-$, CD64$^-$, MHCII$^+$ and CD11c$^+$. C) Quantified frequency of DC subsets isolated from SI LP at 0-14 DPI with *Hpb* or naïve controls. Data shown are pooled from 2 independent experiments (n=5 each) C) Quantified absolute cell number of DC subsets isolated from SI LP at 7DPI with *Hpb* or naïve controls. Data shown are pooled from 2 independent experiments (n=5 each). Quantification of CD40$^+$(E), CD80$^+$(F), CD83$^+$(G), CD301b$^+$(H), PD-L2$^+$(I) within DC subsets isolated from SI LP at 0-14 DPI with *Hpb.* Data shown are from 1 independent experiment (n=5 each). **$p<0.01$, ***$p<0.001$,****$p<0.0001$ as assessed by two-way ANOVA with Šídák's post-test correction for multiple comparisons. ns=not significant.

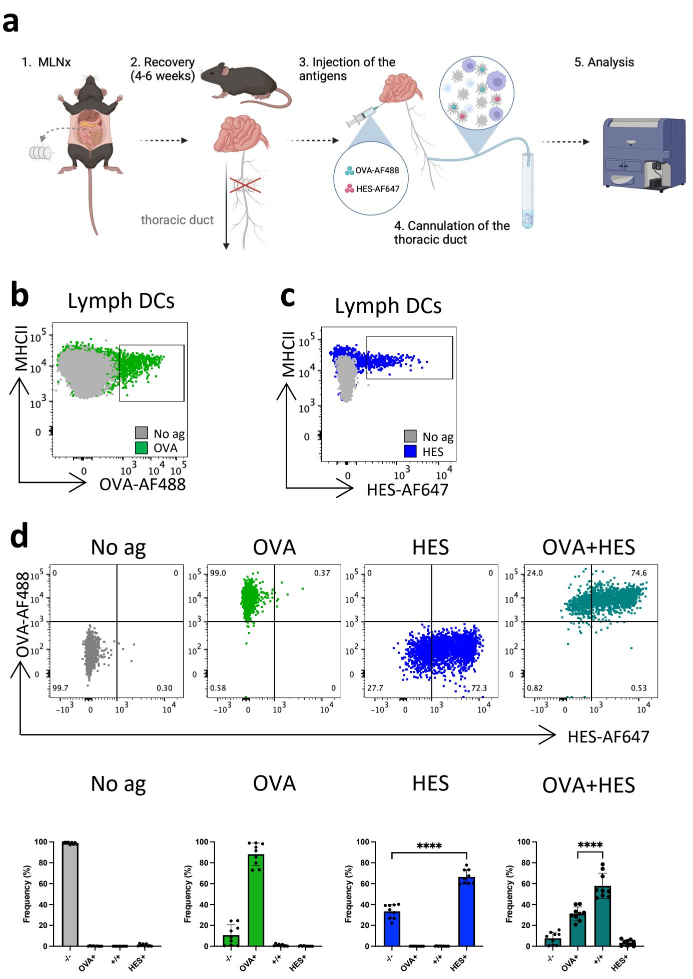


**Figure S2. *Hpb*-derived antigen trafficking and processing by intestinal DCs.**

A) Illustration of lymphadectomy of MLN, and subsequent cannulation of the thorasic duct containing SI LP draining pseudo-afferent lymph. Identification of OVA-AF488 (B) and HES-AF647 (C) carrying DCs in the pseudo-afferent lymph following an injection in the intestinal wall. D) Representative dotplots(top panel) and quantification of proportion (bottom panel) illustrating OVA and HES uptake in OVA-AF488 and/or HES-AF647-pulsed DC subsets isolated from SI LP at steady state.

**a**

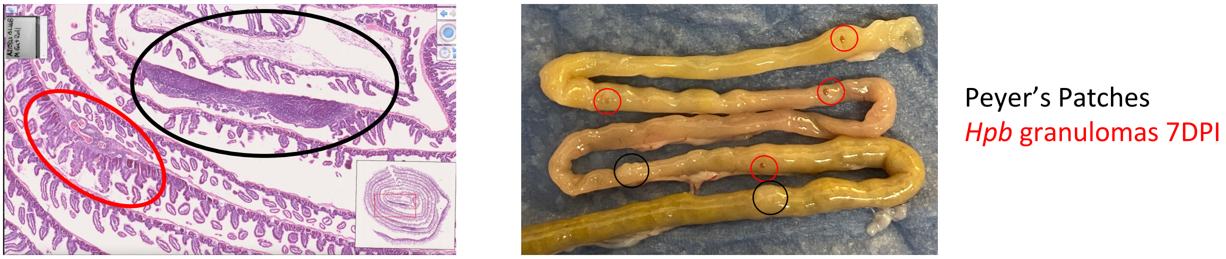


**b**

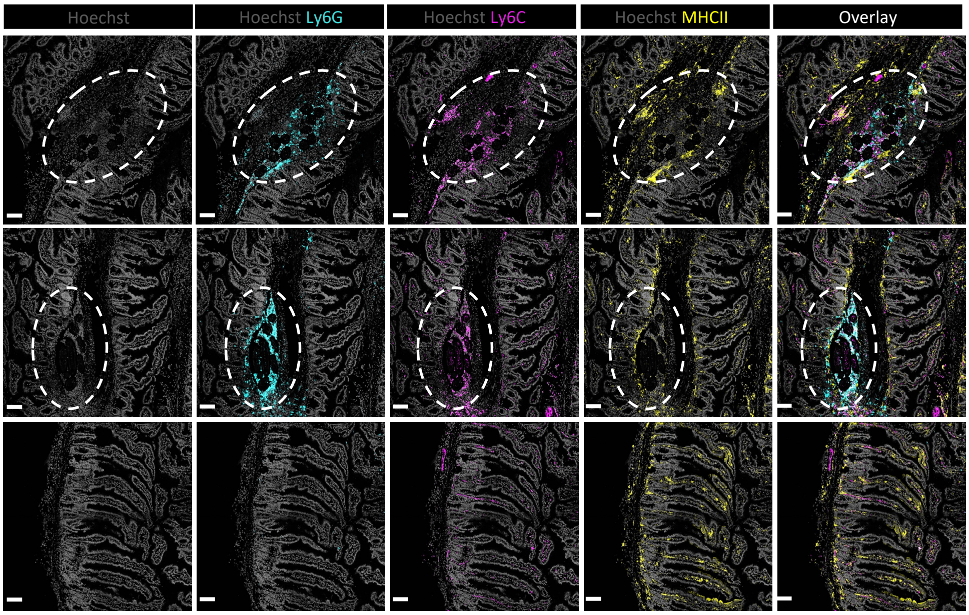


**Figure S3. Visualisation of granulomatous vs. non-granulomatous SI LP tissue during *Hpb* infection.**
A) Visualisation of *Hpb* granuloma (red circles) and Peyer's Patch (black circles) in SI at 7DPI *Hpb*, H&E histology (left) and photograph (right). B) Imaging of granuloma (top, middle panels) and non- granuloma (bottom panel) tissue stained for nuclear (Hoechst), Ly6G, Ly6C and MHCII. Shown as individual stains and then all overlayed.

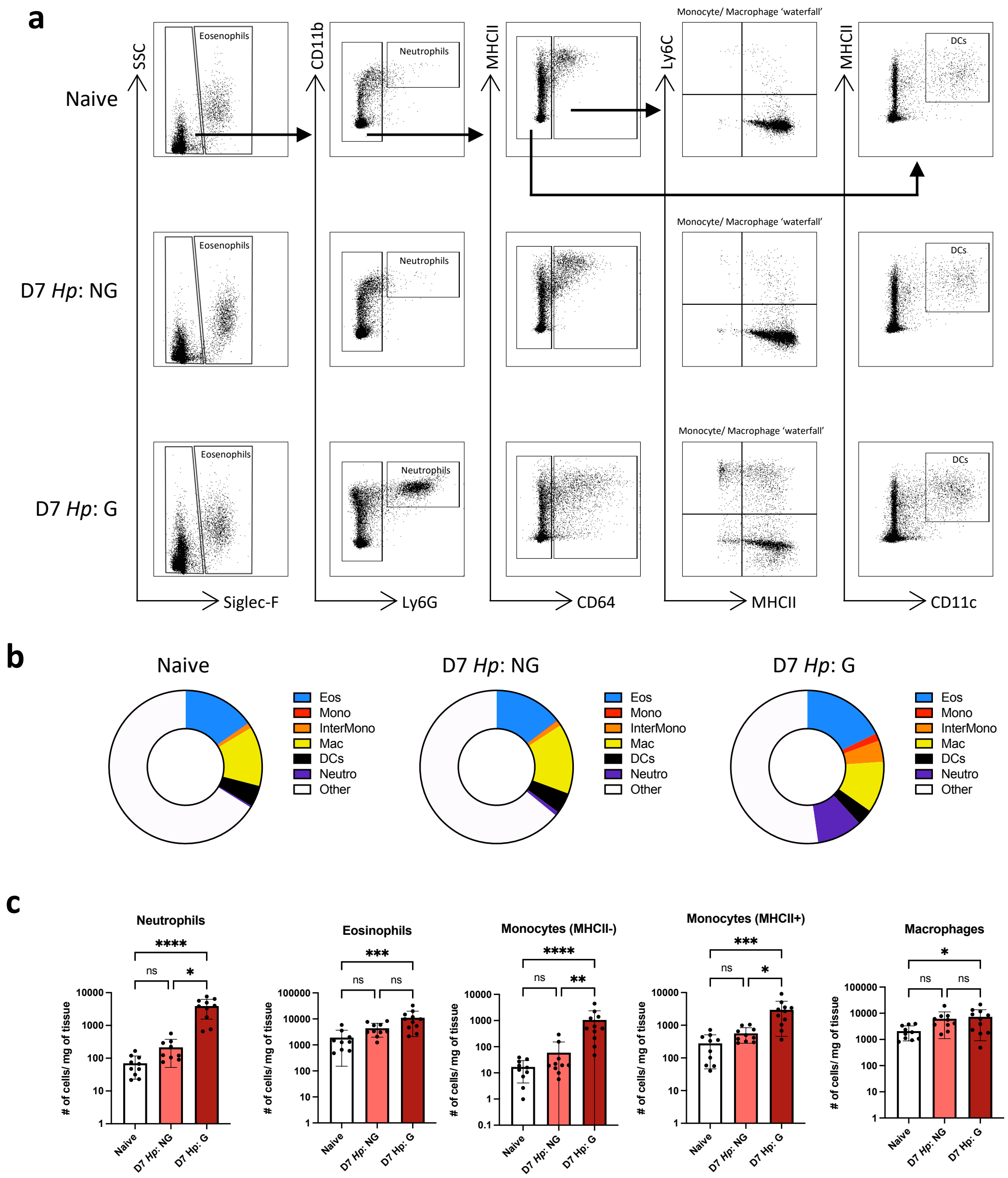


**Figure S4. Characterisation of granulomatous vs. non-granulomatous SI LP tissue during *Hpb* infection.** A) Representative dotplots of gating strategy to for identifying eosinophils (Siglec-F$^{+}$), neutrophils (Siglec-F$^{-}$ Ly6G$^{+}$), monocytes (Siglec-F$^{-}$, CD64$^{+}$, Ly6C$^{+}$, MHCII$^{+/-}$), macrophages (Siglec-F$^{-}$, CD64$^{+}$, Ly6C$^{-}$, MHCII$^{+}$), and DCs (Siglec-F$^{-}$, CD64$^{-}$, CD11c$^{+}$, MHCII$^{+}$) isolated from SI LP at 7DPI with *Hpb* (either granuloma enriched tissue (G) or non-granuloma tissue (NG)) or naïve controls. Pre-gated on live, single, CD45$^{+}$, CD3$^{-}$, B220$^{-}$. B) Proportional distribution of eosinophils, neutrophils, monocytes, macrophages and DCs isolated from SI LP at 7DPI with *Hpb* (either granuloma enriched tissue (G) or non-granuloma tissue (NG)) or naïve controls. C) Quantification of eosinophils, neutrophils, monocytes, macrophages as cells/mg of tissue isolated from SI LP at 7DPI with *Hpb* (either granuloma enriched tissue (G) or non-granuloma tissue (NG)) or naïve controls. **p<0.01, ***p<0.001,****p<0.0001 as assessed by one-way ANOVA (with Dunnett's post-test for multiple comparisons). ns=not significant.
